# Linkage disequilibrium moderates how evolving genetic background shapes variant-level architecture

**DOI:** 10.64898/2026.10.01.756092

**Authors:** Hanbin Lee, Jonathan Terhorst

## Abstract

Evolutionary theory offers a unifying explanation for the genetics of complex traits and diseases by modeling how a trait and its genetic basis are maintained by evolutionary forces. A growing body of evidence suggests that linkage disequilibrium (LD) couples a variant’s evolution to that of the genetic background it resides on. However, widely used single-variant theories omit this coupling between variants and the background, or hold the background fixed. We close this gap by developing a model of linkage LD evolution accounting for stabilizing selection, recombination, and genetic drift. The model deduces the genetic architecture of individual variants subject to the co-evolving genetic background. More efficient selection in larger populations is partially offset by slower decay of existing LD and increased generation of new LD. Conversely, noisier LD in smaller populations exposes the variants to episodes of stronger selection. We ask whether ancestry differences in the relationship between effect-size magnitude and minor allele frequency require different selection strengths or can arise from differences in demographic history alone. Stabilizing selection needs to be more than three times stronger in Africans than in Europeans under the constant-size demography, whereas an Out-of-Africa demography alone can explain most of the observed frequency-dependence gap. As this example indicates, selection strength is in general not identifiable from the frequency–effect-size relationship alone, because different combinations of selection and demographic history can produce the same pattern in data with further redundancy added by LD.

## 1 Introduction

Evolution offers a comprehensive explanation for the patterns of genetic architecture found in genetic association studies. Notably, effect sizes decline with minor allele frequency – a pattern known as frequency-dependence – which is attributed to negative selection (Zeng et al., 2018; O’Connor et al., 2019; Schoech et al., 2019; Zeng et al., 2021). In this model, variants negatively affecting fitness are removed by natural selection, keeping newly emerging deleterious alleles uncommon. This led to studies inferring evolutionary information from frequency-dependence observed in genetic association studies (Zeng et al., 2018; Schoech et al., 2019).

The decline can also be explained by stabilizing selection, which favors individuals with intermediate trait values (Koch et al., 2024; Simons et al., 2025). At the variant level, both mechanisms have the same consequence: variants with larger effects are less common than those with smaller effects (Robertson, 1956). Because numerous genetic variants contribute to complex traits that may affect fitness, the stabilizing-selection framework has proven to be an effective and parsimonious means of explaining complex-trait architecture (Simons et al., 2018; Koch et al., 2024; Patel et al., 2024; Simons et al., 2025; Berg et al., 2025). In such models, heritability is a key parameter controlling the overall contribution of genetics to trait variability, while selection shapes the distribution of effects across variants according to allele frequency, making heritability and frequency-dependence central summaries of genetic architecture. Biobank-scale genotype and phenotype datasets now make it possible to estimate these summaries with increasing accuracy across populations (Gallagher, Ginsburg, and Musick, 2024).

Under stabilizing selection, selection acts on phenotypes and the associated variants inherit the selective pressure from the traits. As a result, selection experienced by each variant depends on the distribution of the other variants constituting the genetic background. This coupling is driven by linkage disequilibrium (LD) between the focal variant and the rest of the variants, because stabilizing selection favors compensatory combinations of alleles rather than acting on each locus independently. The association creates a systematic downshift in genetic variance – known as the Bulmer effect – and further slows selection acting on genetic variants (Bulmer, 1971; Bulmer, 1974; Veller and Coop, 2024; Negm and Veller, 2026). Nevertheless, existing theory and methods have treated variants as evolving independently regardless of the genetic background (Simons et al., 2018; Schoech et al., 2019; O’Connor et al., 2019; Zeng et al., 2018; Zeng et al., 2021; Koch et al., 2024; Simons et al., 2025), or have treated the background as fixed and deterministic in the infinite-population limit (Negm and Veller, 2026; Li, Hermisson, and Sachdeva, 2026). Such limitations obscured the feedback between variants and the evolving genetic background, as well as how changing demography and population size affect frequency-dependent architecture.

To close these gaps, we developed a theory of evolving LD that constitutes the genetic background under all the aforementioned evolutionary forces: stabilizing selection, recombination, genetic drift, and changing demography. The model predicts that, unlike allele frequencies, which change gradually over time, LD – the association between variants – is subject to rapid and sizable random fluctuations that propagate into trait heritability and selection acting on individual variants. Therefore, heritability and frequency-dependent architecture become transient objects subject to temporal change.

We present simulation results isolating these distinct effects of LD in finite populations. When the population expands, slower erosion of existing LD and greater generation of new LD by stabilizing selection buffer the increasing efficacy of selection on variants. Conversely, faster erosion and increasing noise in LD as the population shrinks expose variants more frequently to stronger selection. As a result, various combinations of demographic history and stabilizing selection intensity can yield the same frequency-dependent architecture. Analyzing genetic association study estimates, we found that an Out-of-Africa demography model with similar selection intensity in African and European ancestries can explain the observed frequency-dependence as well as a constant-size demography with selection more than three times stronger in Africans than in Europeans. Consequently, frequency-dependence reflects past selection and demographic history more than just present-day selection.

## 2. Results

### 2.1 Genetic background moderating selection is noisy in finite populations

For an additive trait with genetic and non-genetic components, the variance of the genetic component is called the genetic variance. The genetic component is the sum of the effects of individual loci, so the corresponding variance comes from two separate parts: genic variance and disequilibrium variance. Genic variance is the sum of the individual contributions of the loci, and disequilibrium variance is the covariance term among loci. It is precisely the latter that involves linkage disequilibrium (LD) (Falconer, 1996).

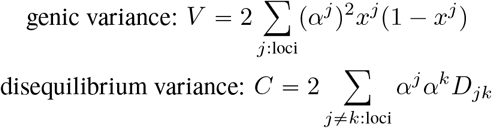

Here, *α*^*j*^ is the effect size of the *j*-th variant, *x*^*j*^ is its allele frequency, and *D*_*jk*_ is the LD covariance between variant *j* and *k*. Throughout the paper, by genetic background we mean genic variance and disequilibrium variance, both adding up to the genetic variance of the trait.

When stabilizing selection acts on traits, the consequence passed down to a locus depends on the genetic background. We refer to *V* and *C* as the genic and disequilibrium variances, respectively. The following formula from Negm and Veller (2026) mathematically formalizes the phenomenon. The selection coefficient is

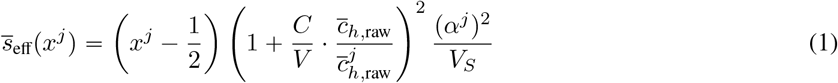

where *x*^*j*^ is the allele frequency and *V*_*S*_ is the selection width of stabilizing selection (the smaller the *V*_*S*_, the stronger the selection). 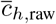 and 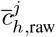 are global and local harmonic means of recombination fractions, respectively, defined as

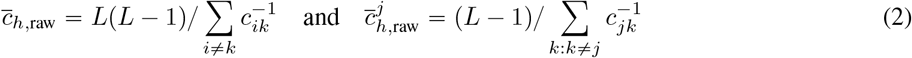

where *L* is the number of loci and *c*_*ij*_ is the recombination fraction between loci *i* and *j*. The “raw” label is to distinguish it from equation (4) that accounts for genetic drift.

It is clear from the formula that selection depends not only on the properties of the *j*-th locus, such as *x*^*j*^ and *α*^*j*^, but also on the genetic background *C* and *V* . The extent to which it depends on *C* and *V* is further controlled by the harmonic recombination fraction 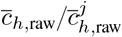. The recombination term enters the formula because the influence of *C* and *V* on selection *s*_eff_ acts through linkage, and stronger linkage (larger 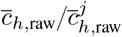) increases this influence. In general, *C* is negative under stabilizing selection, so 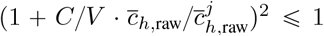 and selection is weakened. Following Negm and Veller (2026), we call this selection reduction Bulmer attenuation to align with the Bulmer effect, which refers to the reduction in genetic variance relative to genic variance caused by negative disequilibrium variance *C* < 0 (Bulmer, 1971; Bulmer, 1974).

The formula rests on two assumptions. First, the backgrounds *V* and *C* are held fixed. This is justified in large populations over short periods, where genic variance changes little (Walsh and Lynch, 2018) and fluctuations in *C* are small (Avery and Hill, 1977). Accordingly, Negm and Veller (2026) tested their theory at *N*_*e*_ *=* 10, 000 over at most *N*_*e*_ */* 10 generations. Second, the recombination rates *c*_*ij*_ in the formula, which control how quickly LD decays, do not account for genetic drift. If drift is omitted, the formula overpredicts the Bulmer attenuation because in reality, genetic drift breaks LD faster than predicted by recombination alone. Since the background’s effect on selection grows with LD, faster erosion of LD will suppress its influence on selection. To this end, we developed a theory that extends Negm and Veller (2026)’s result to the long-term evolution of small populations in which the background evolves and fluctuates.

Before doing so, it is helpful to see how variable the backgrounds *V* and *C* are in neutral populations where selection is absent. In a neutrally evolving population, the expected LD between any pair of loci is zero, but associations between loci can arise spontaneously in finite populations from sampling (Hill and Robertson, 1968; Ohta and Kimura, 1971; Song and Song, 2007). Because LD contributes through pairs of loci, many small associations can add up across a highly polygenic trait. Avery and Hill (1977) showed that the resulting between-population variability of additive variance is mostly due to randomly fluctuating LD rather than allele frequency changes at individual loci.

Figure 1A illustrates this behavior using 100 replicate forward simulations of a neutral polygenic trait in diploid populations of size *N*_*e*_ = 500 over 2,000 generations. The left panel shows the mean trajectories of genic variance (blue) and total additive genetic variance (orange), with bands indicating one standard deviation across replicates. Their means are nearly identical, but total additive variance has a greater spread because it includes the contribution of LD. The middle panel compares their terminal distributions through a quantile-quantile plot: low quantiles of total additive variance fall below the *y* = *x* line, while high quantiles rise above it.

**Figure 1.**
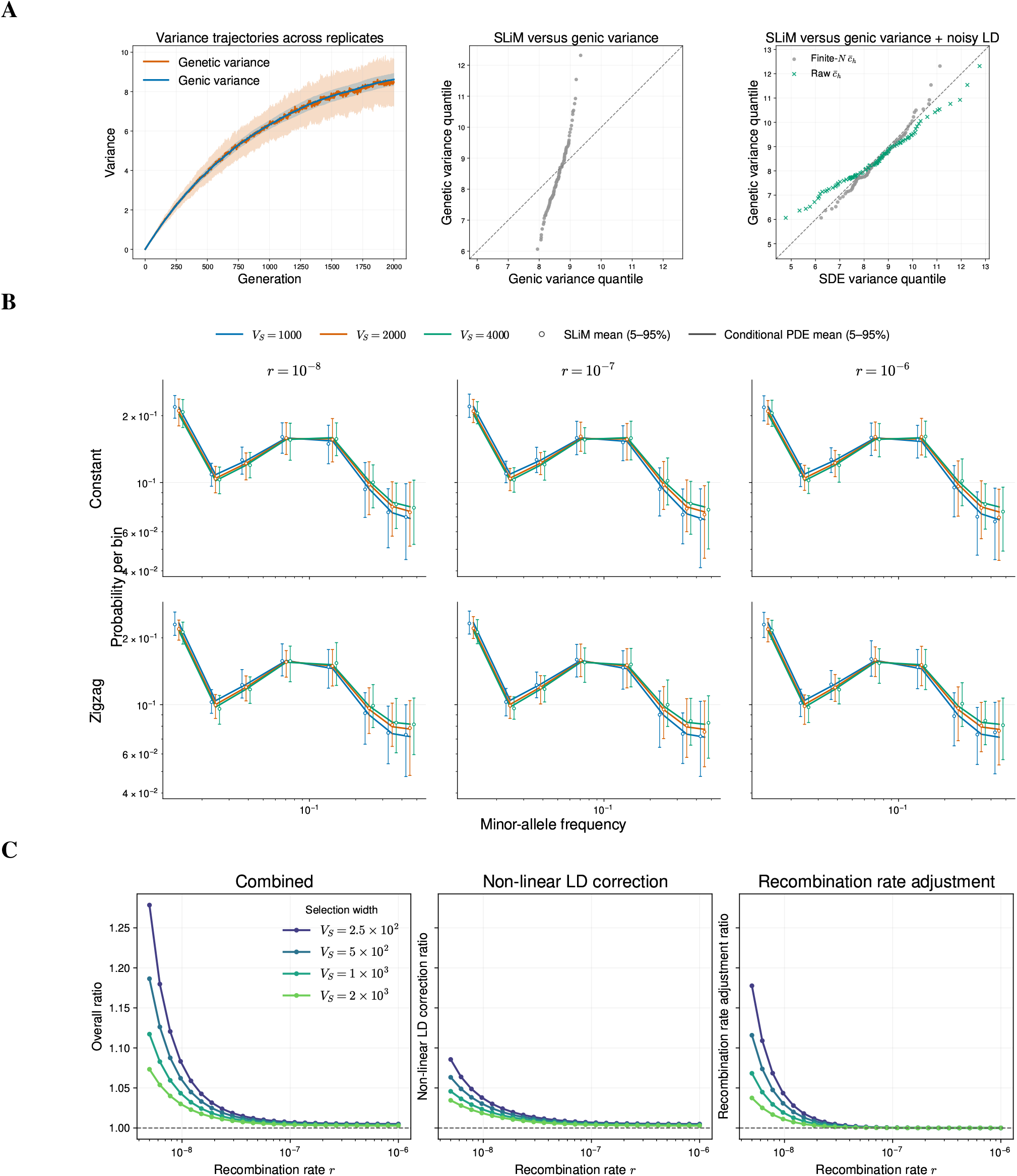
The stochastic model reproduces population- and locus-level dynamics of individual-based simulations. **A** Random changes in additive genetic variance under finite-population sampling. Genetic and genic variance across 100 replicate neutral simulations of a polygenic trait over 2,000 generations in diploid populations of size *N*_*e*_ = 500. We used mutation rate *u* = 5 × 10^−9^, recombination rate *r* = 1 × 10^−8^, genome length *L* = 1 × 10^8^, and mutational variance 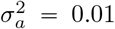. The left panel shows mean trajectories, with bands indicating one standard deviation across replicates. The middle panel compares terminal quantiles of total additive and genic variance, and the right panel compares total additive variance with genic variance plus the stochastic LD contribution from equation (3). **B** The folded sample frequency spectrum. Columns show recombination rates *r* = 10^−8^, 10^−7^, and 10^−6^, and rows show constant and Zigzag demographies. Both rows use mutation rate *u* = 10^−10^ and mutational variance 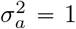. Open circles and error bars show the SLiM mean and 5–95% interval, while solid curves and bands show the corresponding SDE prediction, for the indicated values of *V*_*S*_. **C** Genetic drift weakens Bulmer attenuation and strengthens locus-level selection. The left panel shows the ratio (8) of the average intensity under the full stochastic setting in equation (20) to that under the deterministic system in equation (19); the middle and right panels show its non-linear LD-correction and finite-population recombination-adjustment factors, respectively. The product of the middle and right ratios equals the ratio in the left panel. Results are shown for *N*_*e*_ = 500, genome length *L* = 10^8^, and the indicated selection widths and recombination rates.

To account for LD’s contribution, we derive a stochastic model of disequilibrium variance *C*. Let *C*_*t*_ be the disequilibrium variance due to (*cis*-)LD of the population after random mating and *V*_*t*_ be the genic variance. Both are indexed by *t* to represent their time-varying nature. We show that the dynamics of *C*_*t*_ can be approximated by a discrete-time stochastic process with mean-reverting behavior (Materials and Methods):

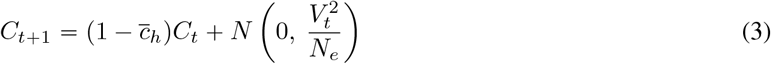

where

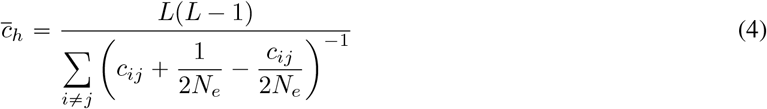

is the harmonic mean of the finite-population-adjusted recombination fractions for all pairs of loci. The finite-population adjustment accounts for the decay of LD due to genetic drift, which degrades LD even when recombination is absent. One could therefore say that this quantity is the effective rate of LD decay that combines recombination and genetic drift. Notably, it prevents extremely small *c*_*ij*_ outliers from exaggerating the outlier-sensitive harmonic mean 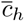, which can be seen by comparing equations (2) and (4). Even if recombination *c*_*ij*_ → 0, the finite 1 / *N*_*e*_ factor keeps *c*_*h*_ above 1 *N*_*e*_ > 0 whereas 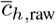 hits 0.

In the equation, existing LD (*C*_*t*_) is discounted by recombination and genetic drift through 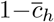, but it is also subject to random noise that grows with genic variance *V*_*t*_. Avery and Hill (1977) derived moment recursions for the random variables derived from *C*_*t*_ and *V*_*t*_, including the LD correlation 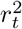 (this is the usual *r*^2^ LD coefficient further indexed by time). Here, we directly model *C*_*t*_ as a stochastic process driven by recombination and drift, with a noise scale that depends on *V*_*t*_, instead of studying its moments. This formulation allows us to couple LD to changing genic variance and, below, extend the model to mutation, stabilizing selection, and non-equilibrium demography. The right panel of Figure 1A compares the simulated total additive variance with *V*_*t*_ *C*_*t*_, where *C*_*t*_ is generated by equation (3). Their close agreement shows that the stochastic LD term accounts for the additional variation missing from genic variance alone. However, using the raw recombination fraction 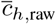 overestimates the spread of disequilibrium variance. This happens because the unadjusted small 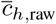 enlarges the LD accumulation, exaggerating the spread.

In sum, additive variance has two distinct components: genic variance that changes smoothly over time and disequilibrium variance, which accounts for most of the temporal noise in additive variance. Since we have seen in the simpler case of neutral evolution that (1) *V* and *C* exhibit noticeable variability and (2) genetic drift erodes LD more quickly than recombination alone, we analyze their impact on selection in the following section.

### 2.2 A model of evolving genetic background reproduces individual-based simulations

We now let the background *V* and *C* evolve under stabilizing selection, along with the allele frequency *x*^*j*^. The evolution of such time-varying random variables is well described in terms of stochastic differential equations (SDEs) (Ewens, 2004; Durrett, 2008). SDEs describe changes in random variables over infinitesimally short intervals through deterministic (*dt*) and stochastic (*dW*_*t*_) components. Readers unfamiliar with SDEs may skip the equations and consult the simulation results, which show their accuracy. Nonetheless, the change from the earlier selection coefficient (1) of Negm and Veller (2026) is operationally minimal: the SDE corresponding to the locus’s allele frequency 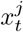 is

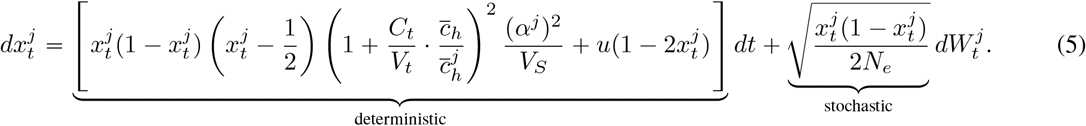

One can see that equation (1) appears in the deterministic term with a few changes. The fixed backgrounds are now indexed by *t*, which means that they evolve over time. The recombination terms 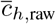 were replaced by drift-adjusted forms 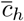. The stochastic term captures random noise due to genetic drift. The symmetric mutation 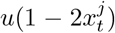 also appears. One might notice the 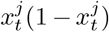 in front of the selection coefficient. This prevents selection from pushing 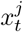 out of the valid range [0, 1] and is standard in diffusion theory (Ewens, 2004; Durrett, 2008).

The backgrounds are no longer fixed inputs and evolve according to

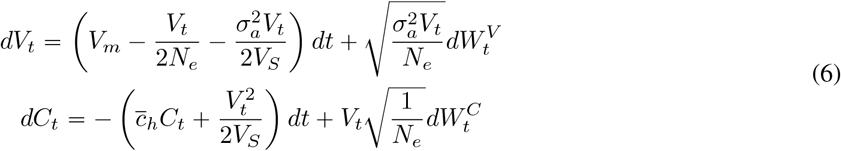

where *V*_*m*_ is the contribution from mutation to genic variance at each generation (see Materials and Methods). The biological interpretation of each term is presented in Table 1. The first equation is standard and has been derived earlier (Barton, 1986; Burger, 2000). We refer interested readers to Appendix A for a more detailed historical exposition. The disequilibrium variance equation for *C*_*t*_ was derived in this work and depends on the genic variance *V*_*t*_. It is critical to assume that *V*_*S*_ ≫ *V*_*t*_ + *C*_*t*_ in our derivation. We later show that this approximation works well in human data, and the mathematical details are in Materials and Methods. It is worth noting that the *C*_*t*_ SDE is a modification of the neutral formula (3) in two respects: the term 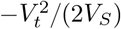 represents new disequilibrium generated by stabilizing selection, which is absent in the neutral setting, and the discrete-generation convention (*t* → *t* + 1) was replaced by a continuous-time representation.

**Table 1:** Description of terms in the deterministic (*dt*) coefficients. See Appendix B for more details.

| Term | Interpretation |
| --- | --- |
| $-\frac{V_t+C_t}{V_S}(m_t - \mu)$ | Selection pulls the mean toward the optimum |
| $V_m$ | New variance input from mutation per generation |
| $-\frac{V_t}{2N_e}$ | Loss of genic variance due to genetic drift |
| $-\frac{\sigma_a^2 V_t}{2V_S}$ | Loss of genic variance due to stabilizing selection |
| $-\bar{c}_h C_t$ | Loss of linkage disequilibrium due to recombination and finite-population sampling |
| $-\frac{V_t^2}{2V_S}$ | New linkage disequilibrium due to stabilizing selection |

We defer the derivations to Materials and Methods, but assess the SDEs’ accuracy against forward simulations. We used two simulations: one in a constant-size population of *N*_*e*_ = 1,000 and another under a ZigZag demography ranging from *N*_*e*_ = 500 to 2,000. The first case shown in Supplementary Figures A1, A2, and A3 demonstrates that the distributions of population statistics *m*_*t*_, *V*_*t*_, and *C*_*t*_ from SDE simulations align well with those from SLiM simulations across recombination rates *r* = 10^−8^, 10^−7^, and 10^−6^, respectively. This agreement also holds outside of constant demography: under a Zigzag history with three expansions and two declines, the SDE reproduces the corresponding terminal distributions from SLiM (Supplementary Figures A4, A5, and A6). The agreement extends from these population-level summaries to the locus-level allele-frequency distribution. Figure 1B compares the folded sample frequency spectra produced by SLiM and by the Kolmogorov equations associated with equation (5). Across the three recombination rates and both the constant and Zigzag demographies, the SDE reproduces the spectra from SLiM over the range of selection widths considered.

It is customary to analyze the equilibrium properties of the genetic variables because the relative contributions of evolutionary forces are more easily seen than under nonequilibrium conditions. In biological terms, equilibrium is reached when evolutionary forces that remove and create the genetic variables eventually balance out, reaching a point where the variables remain unchanged. Nonetheless, we are working in a finite-population setting where genetic drift consistently introduces noise (through stochastic *dW*_*t*_ terms in the SDEs), so the balance is an unchanging distribution rather than a fixed value. This balanced distribution is called the stationary distribution (or law). The genic variance has a Gamma stationary distribution, the LD process has a negative stationary mean, and the mean process has the same stationary law as the Ornstein-Uhlenbeck process (equations 15 and 16; Materials and Methods).

We can use the stationary formula to characterize how genetic drift moves the stochastic system of equations (5) and (6) away from the deterministic formulation of equation (1). As mentioned earlier, the departure from the deterministic theory is driven by two distinct mechanisms: faster LD erosion by drift and noise in the background that modulates selection.

### 2.3 Genetic drift weakens the Bulmer attenuation of stabilizing selection

In the deterministic theory of infinite populations, the fixed background variables *V* and *C* in Negm and Veller (2026) were given numbers, not necessarily derived from the evolutionary configuration of a population. The background equations (6) relax the premise: the background values automatically emerge from the evolutionary configuration, thereby providing the right numbers for *V* and *C* under the deterministic setting. As stated earlier, we work under the equilibrium setting. When equilibrium is reached, the distribution of the background remains constant. This means that the expectations of *V*_*t*_ and *C*_*t*_ no longer depend on *t*, so it is natural to replace the fixed *V* and *C* of equation (1) with their respective expectations. Then, we have

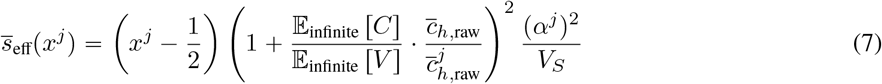

where the expectation is computed with respect to the stationary distribution of the evolutionary path, using 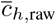 instead of the adjusted 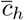. Note that the expectation E [·] is also implicitly a function of 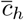, separately from the 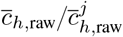 because the stationary law depends on the recombination fraction. That is, E _finite_ [·] and E _infinite_ [·] use 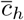 and 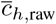, respectively.

The Bulmer attenuation involving the background is our main focus. We can deduce from equation (5) that this quantity in finite populations is the expectation of the squared ratio

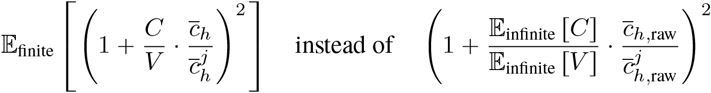

which is the squared ratio of expectations under the polygenic assumption (Materials and Methods). For a cleaner comparison between the two, we omit the ratio 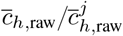 and observe the following decomposition:

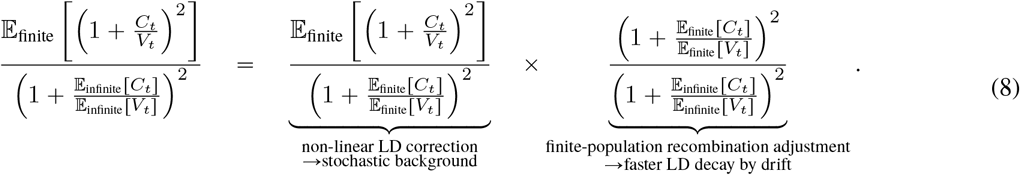

The decomposition reflects the aforementioned two mechanisms, and we explain the biological and mathematical principles behind them.

The first factor comes from the fluctuating background. By equation (5), selection on a variant is stronger when *C*_*t*_ *V*_*t*_ is less negative than usual and weaker when it is more negative. Because the attenuation 1 + *C*_*t*_ */ V*_*t*_ ^2^ is quadratic, these two departures are not symmetric: a shift toward zero raises selection more than an equal shift away from zero lowers it, so over time the gains during weak-LD periods exceed the losses during strong-LD periods. Thus, the average Bulmer attenuation is smaller than the attenuation at the average background. The second factor comes from the LD decay rates 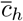 versus 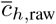. The finite-population 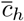 is larger than 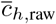, especially in smaller populations, because genetic drift erodes LD in addition to recombination (Equation 4).

In the simulations presented in Figure 1C, we measured these effects in a population of *N*_*e*_ = 500. The middle panel shows that the non-linear LD correction increases selection by up to approximately 9%, while the right panel shows that the finite-population recombination adjustment increases it by up to approximately 18%. Both effects are strongest when recombination is weak and stabilizing selection is strong (small *V*_*S*_), because these conditions generate more negative LD and therefore leave more scope for drift to weaken the Bulmer attenuation. Their product, shown in the left panel, raises the effective selection intensity by approximately 28% relative to the deterministic, infinite-population prediction. All three ratios approach one as recombination increases, because rapidly decaying LD contributes little attenuation in either model.

As the Bulmer attenuation reduces selection experienced by variants, Negm and Veller (2026) suggest revising the raw selection intensity inferred from genetic association studies upward. Our results imply that the required revision is smaller than the deterministic prediction, more so in smaller populations, because genetic drift reduces the attenuation.

### 2.4 Bulmer attenuation weakens frequency-dependent architecture in equilibrium

We next ask how these locus-level dynamics shape the distribution of effect sizes across allele frequencies. It is common to summarize genetic architecture by asking whether rare alleles tend to have larger effects than common alleles. Such frequency-dependence is usually interpreted as a signature of selection. Specifically, genome-wide association study (GWAS) estimates exhibit a negative correlation between heterozygosity *x*^*j*^ ( 1 − *x*^*j*^ )*x*^*j*^ is the allele frequency) and effect size *α*^*j*^ across loci, a pattern commonly referred to as frequency-dependent architecture.

Lee and Terhorst (2026) derived a formula for the expected squared effect size E [(*α*^*j*^) ^2^ | *x*^*j*^ (1 − *x*^*j*^)] as a function of heterozygosity, under a strong-recombination assumption that largely ignores selection-induced LD

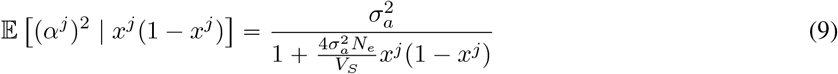

at the stationary equilibrium. They used an evolutionary linear mixed model (LMM) to confirm this formula in individual-based forward simulations (Haller, Ralph, and Messer, 2025). The SDE approximation of equation (5) offers a faster simulation scheme that allows us to use data-driven methods to learn the frequency-dependent architecture.

Using a large number of (*α*^*j*^, *x*^*j*^) pairs simulated with SDE (5), we fitted non-increasing spline curves to estimate E [(*α*^*j*^) ^2^ | *x*^*j*^ (1 − *x*^*j*^)] under constant demography with sufficiently long burn-in to ensure equilibrium. In Figure 2A, the data-driven spline curves closely resemble the approximate formula (9) across varying selection intensities. For lower recombination rates than those explored in Lee and Terhorst (2026), the gap between the theoretical (green) and data-driven (red) curves widens (Figure 2B). In contrast, our stochastic model’s prediction (purple), which adjusts for the Bulmer attenuation,

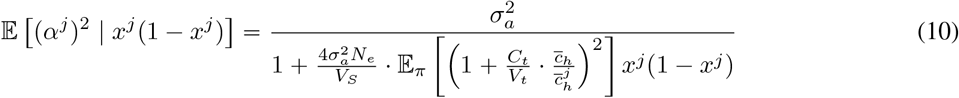

remains close to the data-driven curve. The attenuation factor 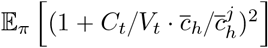 is less than 1, so squared effect sizes drop more slowly in equation (10) than in (9). In the figure, we assumed 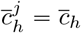.

**Figure 2.**
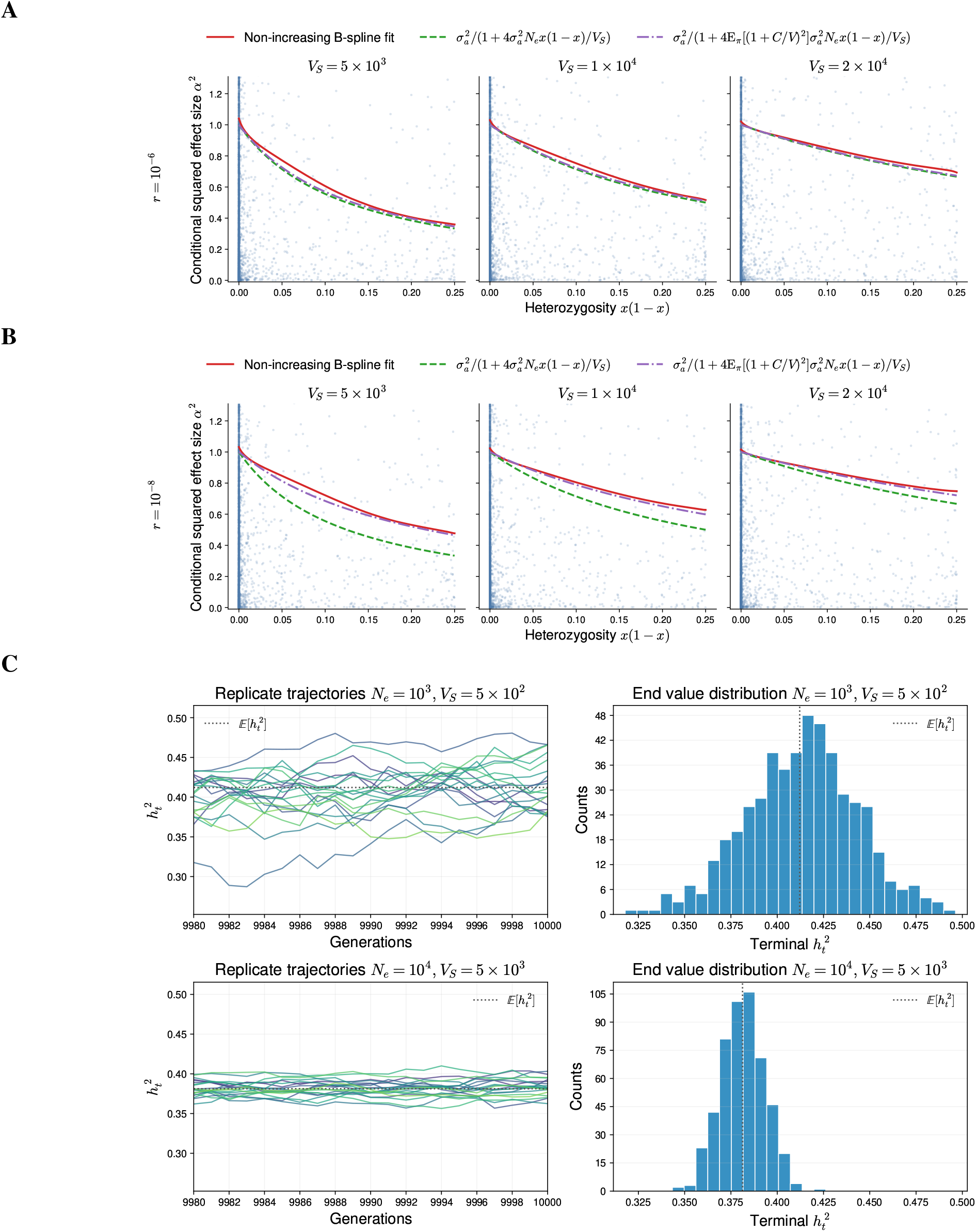
Frequency-dependent architecture and heritability under the stationary equilibrium. **A–B** Frequency-dependent architecture of allelic effect sizes under stabilizing selection. The simulations were run under constant *N*_*e*_ 10, 000 demography and *L* = 10^8^ with recombination rates *r* = 10^−6^ (**A**) and 10^−8^ (**B**), and varying *V*_*S*_ values. The green dotted curves represent equation (9) and the purple dotted curves represent equation (10). **C** Heritability remains variable at stationary equilibrium. Narrow-sense heritability 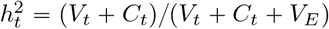 was simulated under constant population size for 10,000 generations. The left column shows the final 20 generations of 20 independent trajectories, and the right column shows the distribution of terminal 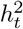 across 500 additional replicates. Rows correspond to (*N*_*e*_, *V*_*S*_) = (10^3^, 5 × 10^2^) and (10^4^, 5 × 10^3^), respectively. Other parameters were genome length *L* = 10^8^, mutation rate *u* = 5 × 10^−9^, recombination rate *r* = 10^−8^, and mutational variance 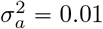. Dotted lines mark 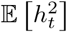 from equation (11).

Since the relative reduction in genetic variance *C*_*t*_ */ V*_*t*_ is larger when linkage is stronger, the formula shows that stronger linkage compromises the correlation between variant effect size and frequency. As shown in Figures 2A and 2B, green and purple lines are almost identical when linkage is weak (high recombination rate, *r* = 10^−6^), while the gap is wider at a low recombination rate (*r =* 10^−8^). The figure also shows that squared effect sizes drop faster when selection is stronger (*V*_*S*_ is smaller), consistent with the intuition that frequency-dependence is a result of selection. However, we will later show that this observation breaks down when demography is no longer constant.

Standard *α*-model parameterizations place heritability explicitly in the squared effect-size formula, as the scale that normalizes per-variant contributions to the total genetic variance (Speed et al., 2017; Schoech et al., 2019). Equation (10) contains no such terms: its scale is controlled by mutational variance 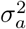, and its shape is set by *N*_*e*_, *V*_*S*_, and the Bulmer attenuation. Heritability must therefore enter differently. We have so far used the term informally, as the share of phenotypic variance that is genetic. The next section defines it within the model as a quantity built from *V*_*t*_ and *C*_*t*_ and shows why it cannot be inserted directly into the formula: it emerges, together with the frequency-dependence curve, from the same evolutionary parameters.

### 2.5 Heritability emerges jointly with frequency-dependent architecture

Heritability is absent from the squared effect-size formula in the preceding section. Readers familiar with the conventional frequency-dependence formula

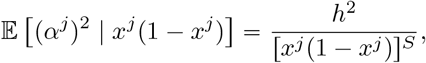

from which heritability *h*^2^ can be directly read off from the numerator (Speed et al., 2017), may find this result confusing. Instead, heritability and frequency-dependence emerge jointly from evolutionary processes. We show that heritability demands an extra disequilibrium component that is missing in the frequency-dependence formula covering only the genic variance.

Narrow-sense heritability is defined as the ratio of additive genetic variance to total phenotypic variance (Falconer, 1996; Walsh and Lynch, 2018). In our model, additive variance is *V*_*t*_+ *C*_*t*_, the sum of genic and disequilibrium variance, while total phenotypic variance is *V*_*t*_ + *C*_*t*_ + *V*_*E*_, where *V*_*E*_ is the non-genetic variance component. Thus,

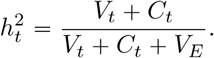

Although *V*_*E*_ is fixed in our model, the additive variance *V*_*t*_ *C*_*t*_ changes over time as genic variance and LD fluctuate. The numerator therefore varies, as does the total phenotypic variance in the denominator. Consequently, *h*^2^ is itself a time-varying random quantity. In particular, even at equilibrium, *h*^2^ has a stationary *distribution* rather than a single deterministic value (Figure 2C).

One common reduction is to report mean heritability, defined as (Yang et al., 2015; Simons et al., 2018)

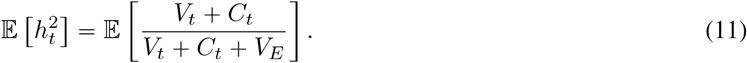

However, this quantity is conceptually different from the heritability of an actual population: it is the average across repeated evolutionary histories generated under the same evolutionary parameters and demographic history (Figure 2C). Another approximation averages the random numerator and denominator separately, yielding

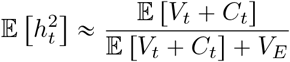

as in methods that assume constant genetic variance (Yang et al., 2015; Gazal et al., 2017; Simons et al., 2018).

Only in special cases (e.g., constant population size and stationarity) can stationary expectations replace time-dependent quantities. Under these additional assumptions, the expected additive variance can be written in terms of the evolutionary parameters of the model, as shown in equations 15 and 16 (see Materials and Methods):

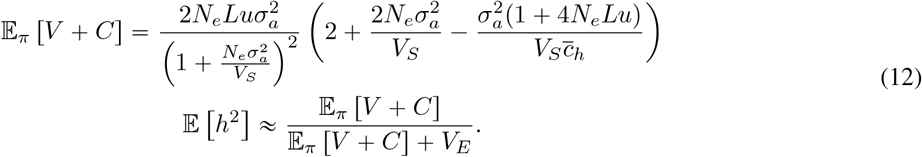

Comparing equations (12) and (10) side by side, we see only the parameters common to the two formulas. No simple transformation from one formula to the other is available. More precisely, heritability estimation sums the contributions of individual variants, (*α*^*j*^) ^2^*x*^*j*^ (1 − *x*^*j*^ ), across the genome (Falconer, 1996; Yang et al., 2015), and evolutionary models compute expected heritability in the same way, by adding up the expected contribution of each variant under free recombination (Keightley and Hill, 1989; Simons et al., 2018; Lee and Terhorst, 2026). Such a sum recovers the genic variance *V*, and it is the only part of heritability that the frequency-dependence curve can inform. The disequilibrium variance *C* is a covariance across loci; no sum over individual variants contains it, and it is also the noisiest part. In the *α*-model, heritability enters the effect-size formula only because it is imposed as a normalization on the distribution of effects; here the total genetic variance is a random output of the evolutionary process, so it is not available to be imposed. The two summaries share parameters because both are outcomes of the shared evolutionary process, but neither determines the other.

Equation (12) is useful because it shows explicitly how heritability depends on evolutionary parameters. It also illustrates why heritability is a many-to-one function: the same value of expected heritability E [*h*^2^] can arise from different combinations of effective population size (*N*_*e*_), mutation rate (*u*), mutational effect variance 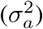, selection strength (*V*_*S*_), and recombination rate 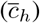. Conversely, the same evolutionary parameters can generate different realized values of 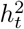 across replicate populations because linkage disequilibrium fluctuates stochastically. Thus, even when expected heritability is known exactly, it does not uniquely identify the underlying architecture.

### 2.6 Linkage buffers demographic change acting on stabilizing selection

The analysis so far has been conditional on equilibrium. Now we study the non-equilibrium conditions under changing demography. Non-equilibrium demography, represented here by changes in population size *N*_*e*_ over time, has three channels affecting selection. The most important route is through reduced genetic drift. Qualitatively, selection becomes more effective as *N*_*e*_ grows. However, the two remaining paths acting through linkage have the opposite effect: as *N*_*e*_ increases, the Bulmer attenuation becomes stronger and reduces selection. One is the increased genic variance *V*_*t*_ in larger populations, which increases the magnitude of *C*_*t*_, and the other is the smaller *N*_*e*_-adjusted LD decay rate 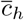, which allows LD to persist longer. These opposing linkage-mediated effects are not strong enough to reverse the increasing efficacy of selection with increasing *N*_*e*_, but they materially temper it.

Consider three populations that constitute the Out-of-Africa (OoA) event modeled in Gutenkunst et al. (2009): the Yoruba (YRI), Utah residents of Northern and Western European ancestry (CEU), and Han Chinese (CHB). The African population’s size remained relatively constant over time, while the latter two populations experienced severe bottlenecks followed by rapid expansion (Figure 3A, first row). The adjusted recombination rate that measures the rate of LD decay immediately responds to the demographic shift (Figure 3A, second row). When *N*_*e*_ declines, 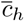 grows and vice versa. Decreasing *N*_*e*_ increases the effects of drift, so LD between closely linked loci breaks more quickly. As a result, the *N*_*e*_-adjusted harmonic mean *c*_*h*_, which is sensitive to small outliers, grows sharply. This effect is present in both Europeans and East Asians, but only when recombination is small (*r* = 10^−8^). When recombination is already strong (*r* = 10^−6^), genetic drift contributes little to the already rapidly eroding LD driven by recombination.

**Figure 3.**
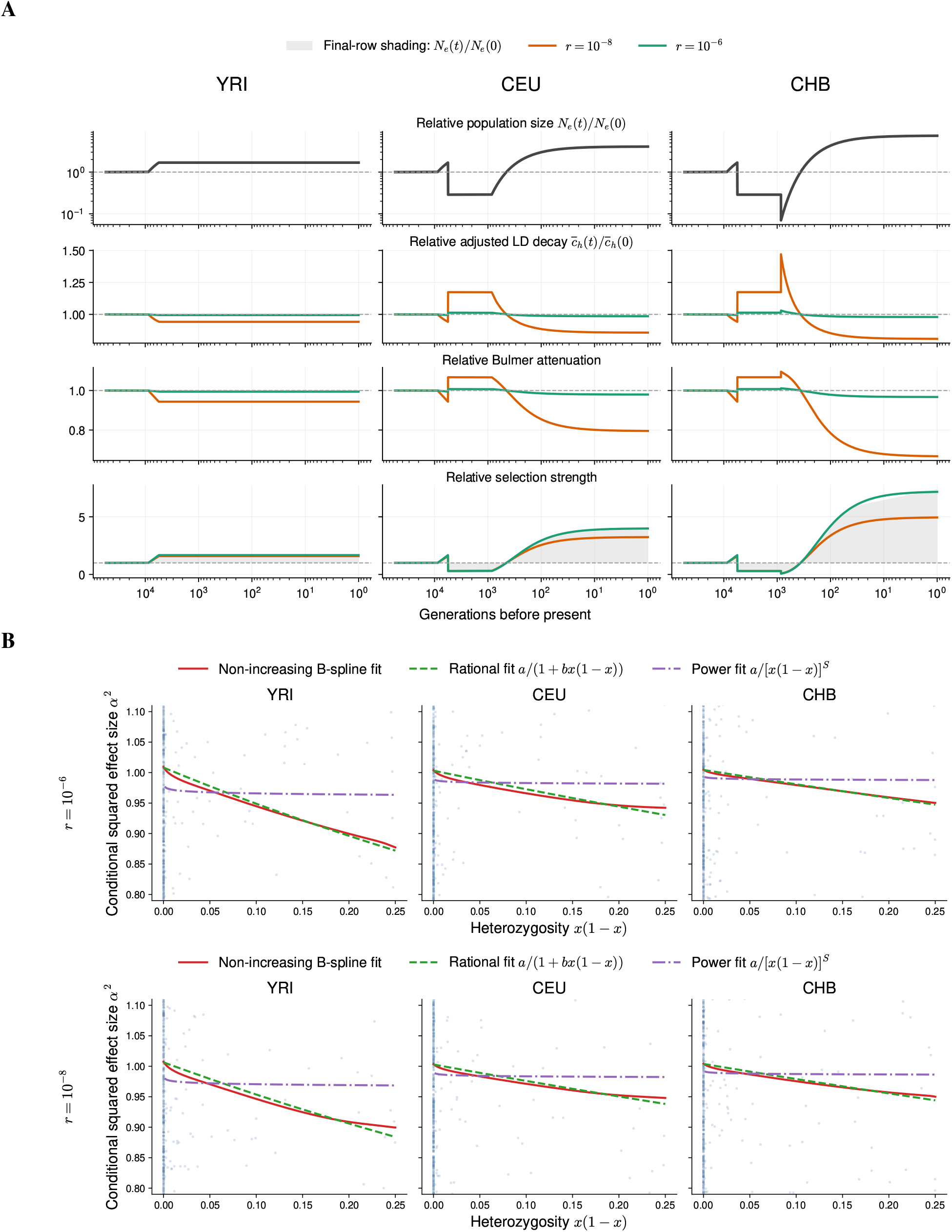
Linkage buffers demographic change and reshapes frequency-dependent architecture. Results use the three-population Out-of-Africa model of Gutenkunst et al. (2009), implemented as `stdpopsim`‘s `OutOfAfrica_3G09`, for Yoruba (YRI), Utah residents with Northern and Western European ancestry (CEU), and Han Chinese in Beijing (CHB). **A** Demography and its consequences for LD-mediated selection through time, with columns corresponding to YRI, CEU, and CHB. From top to bottom, rows show effective population size *N*_*e*_ *(t)*, the finite-population-adjusted LD decay rate 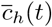, Bulmer attenuation, and the resulting strength of selection, each relative to its ancestral value. Black curves show relative population size; orange and green curves show results for recombination rates *r =* 10^−8^ and 10^−6^, respectively. Shading in the bottom row reproduces relative population size to show the gap between demography and selection. **B** Present-day frequency-dependent architecture for YRI, CEU, and CHB at *V*_*S*_ 50,000 and genome length *L =* 10^8^. The upper and lower rows use *r =* 10^−6^ and 10^−8^, respectively. Curves show a non-increasing B-spline fit, the reciprocal fit *a / [*1 + *bx (*1 − *x)* with *a, b* ⩾ 0, and the power-law *α*-model fit *a / [x* + ( 1 − *x*)] ^*S*^ with *a, S* 0. Present-day selection is strongest in CHB, although its frequency-dependence is weakest.

Once the LD decay rate increases, the Bulmer attenuation (1 + *C*_*t*_ */ V*_*t*_ )^2^ follows, but in a more gradual manner because it takes time for the negative *C*_*t*_ to decay according to the newly increased rate (Figure 3A, third row). Therefore, right after the bottleneck, selection does decrease, but the simultaneous reduction in the Bulmer attenuation partially buffers it. The same mechanism runs in reverse during expansion: 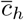 falls, the Bulmer attenuation strengthens, and it now works against the rising efficacy of selection. Once the populations start to expand, selection closely tracks the population size. Nevertheless, when recombination is weak (*r =* 10^−8^), selection lags behind the population growth (Figure 3A, fourth row). This is precisely due to the Bulmer attenuation presented in the third row, which again stems from the reduction in LD decay rate 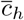 as *N*_*e*_ grows.

### 2.7 Demographic history reshapes frequency-dependent architecture

Natural populations have undergone complex demographic events, exemplified by the human migration Out-of-Africa. In such cases, closed-form equations such as equation (10) from the previous section are not available. While the equations derived from equilibrium conditions predict that stronger selection leads to a steeper decline in effect size with increasing minor allele frequency, this relationship does not hold once demography changes over time. Instead, frequency-dependent architecture depends on the entire history of the population and not just the contemporary strength of selection.

By simulating the locus evolution model (5) to track time-varying *N*_*e*_ under an Out-of-Africa demography (Figure 3A) (Gutenkunst et al., 2009), we find that frequency-dependence is weakened by bottleneck events followed by rapid expansion. In Figure 3B, we see that the average squared effect sizes drop more gradually in two non-African populations than in the Yoruba population. Nevertheless, present-day selection is strongest in CHB, where frequency-dependence is weakest, followed by CEU and YRI in that order. That is, frequency-dependent architecture does not reflect the present-day selection that the populations are experiencing.

This shows that the same selection and recombination parameters can generate different apparent frequency–effect-size relationships across populations with different demographic histories. As speculated earlier by Lee and Terhorst (2026), the quotient function of the form *a / (*1 + *bx* (1 − *x*) approximates E [(*α*^*j*^) ^2^ | *x*^*j*^ (1 − *x*^*j*^)] well, unlike the *α*-model (which controls selection through *S* ⩾ 0 in [*x*^*j*^ 1 − *x*^*j*^] ^−*S*^), which poorly captures the declining trend. However, as shown above, the steepness of the decline in effect-size magnitude combines all the evolutionary forces involved and should not be interpreted as an indicator of selection strength (Zeng et al., 2018; Schoech et al., 2019; Zeng et al., 2021). The present-day curve depends not only on the current evolutionary parameters, but also on the population history through which it was generated.

### 2.8 Present-day frequency-dependence does not uniquely identify evolutionary parameters

To connect these theoretical predictions to empirical estimates, we extracted the meta-analysed conditional squared effect-size estimates and corresponding minor allele frequencies reported by Rossen et al. (2026) (Figure 4A). Our earlier work showed that squared effect-size estimates stratified by frequency approximate frequency-dependence in a curve-fitting framework (Lee and Terhorst, 2026). We considered estimates defined by African-ancestry allele frequency bins among variants common in European ancestry and European-ancestry frequency bins among variants common in African ancestry.

**Figure 4.**
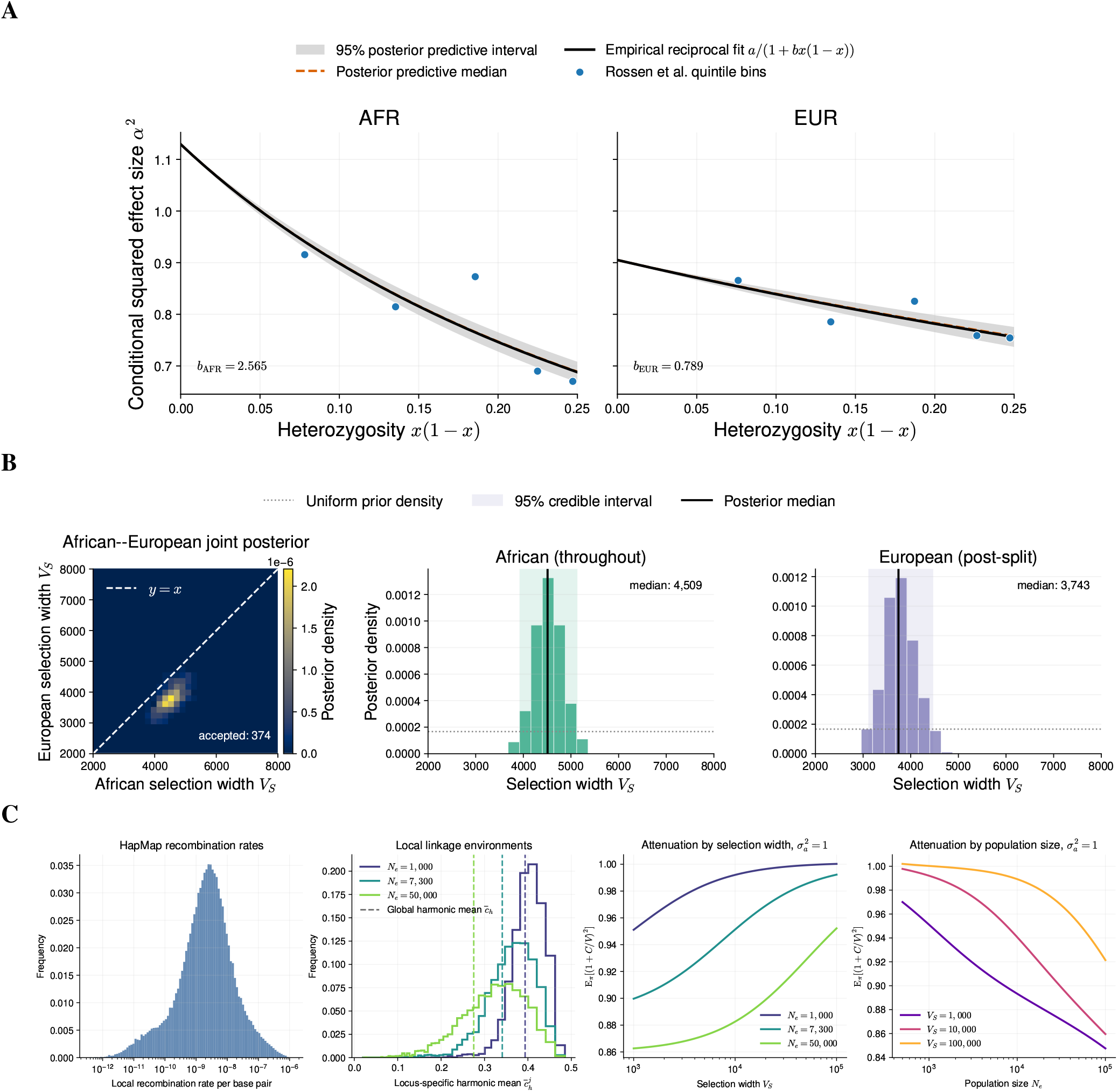
Stronger frequency-dependence in Africans than in Europeans is compatible with similar selection intensity. **A** Empirical and joint posterior-predictive frequency-dependent effect-size curves. Points show the meta-analysed conditional squared effect-size estimates of Rossen et al. (2026) in African-ancestry minor-allele-frequency bins among variants common in European ancestry (left) or European-ancestry bins among variants common in African ancestry (right). Black curves are reciprocal fits *a / [x* + ( 1 − *x*)] to the five empirical bins, with *b*_AFR_ = 2.565 and *b*_EUR_ = 0.789. Orange dashed curves and gray bands show the medians and pointwise 95% intervals from joint posterior-predictive simulations. **B** Joint ABC posterior density for the African selection width, which also applies to the ancestral history, and the post-split European selection width (left), with a white dashed *y = x* line indicating equal values, and their marginal posteriors (center and right). The color scale gives joint posterior density; all panels use the same particles accepted jointly using both empirical coefficients. Black lines and shaded regions give marginal posterior medians and 95% credible intervals, and dotted lines give the marginal prior density. **C** Local linkage properties computed from the HapMap Phase II GRCh37 recombination map. From left to right, the panels show the distribution of local recombination rate; the distribution of locus-specific finite-population-adjusted harmonic mean recombination fractions 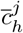 for 3,000 focal loci, with dashed lines giving the corresponding genome-wide harmonic means 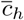 and colors indicating effective population size; stationary Bulmer attenuation E_*π*_ [(1 + *C*/*V* )^2^ ] across selection widths for three effective population sizes; and the attenuation across effective population sizes for *V*_*S*_ = 10^3^, 10^4^, and 10^5^.

The conditional squared effect size declines more sharply when variants are ordered by African rather than European frequency. Fitting the reciprocal curve *a /* [1 + *bx* (1 − *x*)] to each five-bin series gives *b*_AFR_ = 2.565 and *b*_EUR_ = 0.789, a 3.3-fold difference. As a simplified exercise, we plugged these estimates into the stationary frequency-dependence formula (10). We calculated stationary Bulmer attenuation over a grid of *N*_*e*_ and *V*_*S*_ using the harmonic mean 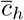 estimated from the HapMap recombination map (Figure 4C). At *N*_*e*_ = 10,000 and 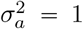, the map gives 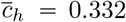. Assuming 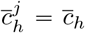, we substituted the grid-specific attenuation into 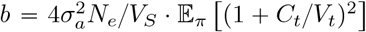 and interpolated the value of *V*_*S*_ matching each empirical *b*. This gives *V*_*S*_ 1.49 × 10^4^ for the African-ancestry curve, with Bulmer attenuation 0.952, and *V*_*S*_ 4.97 × 10^4^ for the European-ancestry curve, with attenuation 0.980. Conversely, fixing *V*_*S*_ and changing *N*_*e*_ can likewise yield the same *b* by following the same steps (Figure 4C). This provides a concrete example of non-identifiability.

We next modeled the shared history while allowing selection strength to change at the population split. We used approximate Bayesian computation (ABC) under the Out-of-Africa demography to infer two selection widths: one shared by the ancestral and African histories, and another for the European history after the split (Figure 4B) (Beaumont, Zhang, and Balding, 2002). The two parameters had independent uniform priors on 2000, 8000, and particles were accepted jointly using both *b*_AFR_ and *b*_EUR_. Among the 374 accepted particles, the posterior medians and 95% credible intervals were 4509 [3932, 5123] for the ancestral and African histories and 3743 [3122, 4460] for the European history. Unlike the three-fold difference in *V*_*S*_ from the stationary back-calculation, the median *V*_*S*_ values inferred for the African and European histories differ by (4509 3743) / 4509 = 17.0%. The joint posterior of selection widths gives the posterior-predictive bands in Figure 4A. At the posterior-median selection widths, the contemporary Bulmer attenuation is 0.917 in Africans and 0.942 in Europeans, corresponding to reductions in effective selection of 8.3% and 5.8%, respectively.

## 3 Discussion

Selection on a complex trait acts on variants through the genetic background on which they reside. We studied the evolution of that background under demographic change: stabilizing selection generates linkage disequilibrium, recombination and drift erode it, and finite-population reproduction continually introduces fluctuations. The resulting theory explains how frequency-dependent architecture and heritability emerge from the combination of these evolutionary forces. In particular, our work focuses on linkage disequilibrium because its dynamics are highly sensitive to genetic drift and demography.

### Stochastic LD links trait- and variant-level architecture

Stabilizing selection on phenotypes creates compensatory associations among alleles (Bulmer, 1971; Bulmer, 1974). Classical quantitative genetic theory describes how such associations alter trait variance, while recent work shows how they attenuate selection experienced by individual variants (Lande, 1975; Veller and Coop, 2024; Negm and Veller, 2026). Our contribution is to let the genic and disequilibrium variances evolve under mutation, selection, recombination, and drift, rather than treating them as fixed inputs to locus dynamics. This extends the deterministic attenuation formula to a distribution of genetic backgrounds and recovers the distributions observed in individual-based simulations under both constant and changing population size.

Two finite-population effects matter for this extension. First, drift changes the background itself: it depletes genic variance and accelerates the decay of LD, reducing the compensatory associations maintained by selection. The adjustment to the harmonic mean of recombination fractions is particularly important for tightly linked pairs, whose LD can decay through drift even when recombination is negligible. Second, selection depends nonlinearly on the fluctuating ratio *C*_*t*_ / *V*_*t*_. Replacing the stochastic attenuation factor by its value at an average background loses a contribution from background variability.

These results modify the upward revision of stabilizing selection proposed by Negm and Veller (2026) for models that omit Bulmer attenuation. Drift weakens that attenuation, so the required revision is smaller than the deterministic prediction, especially in smaller populations with strong linkage. There is consequently no universal multiplicative correction that can be applied to an estimate of selection independently of population size and recombination. Hence, descriptions of genetic architecture (Koch et al., 2024; Simons et al., 2025) must account for this dependence when they are transferred between populations.

### Present-day frequency-dependence is evolutionarily underdetermined

Existing works commonly characterize frequency-dependence using the phenomenological *α*-model (Zeng et al., 2018; Schoech et al., 2019; Zeng et al., 2021; Ashraf and Lawson, 2021; Taylor and Lawson, 2024; Rossen et al., 2026). Because this model is not itself an evolutionary model, its parameters do not directly identify the underlying evolutionary parameters. Some studies instead calibrate the statistical parameters against forward simulations by finding evolutionary configurations that produce estimates similar to those observed empirically. Such calibration assumes that the modern frequency-dependence curve contains enough information to distinguish among the candidate evolutionary models. This work shows why matching this summary alone cannot identify a unique evolutionary configuration. Combinations of different selection intensities and demographic models can yield the same frequency-dependent architecture.

Changes in population size affect architecture through processes with different response times. The rate of losing genetic variation by drift and the drift-adjusted LD decay rate respond immediately to a change in *N*_*e*_, whereas genetic variance and allele frequencies retain information about earlier generations. Expansion increases selection while gradually strengthening the Bulmer attenuation that partially offsets it. A bottleneck has the converse effects, reducing the efficacy of selection while weakening its attenuation by LD. The frequency-dependence curve holds the accumulated records of these processes. In our Out-of-Africa simulations, the population with the strongest present-day population-scaled selection can have the weakest frequency-dependence (Figure 3B).

### Reporting heritability and genetic architecture

Heritability and frequency-dependence reflect different aspects of the same genetic system. The frequency-dependence curve describes single-locus effects conditional on frequency; combined with the distribution and number of loci, it informs genic variance. Additive genetic variance also includes the covariance across loci, the disequilibrium variance *C*_*t*_. Consequently, neither a frequency-dependence curve nor a heritability value determines the other.

Summing single-locus variance contributions recovers *V*_*t*_, whereas the variance of a realized genetic value includes LD covariances. In human statistical genetics, genomic mixed models such as GCTA estimate the variance explained collectively by typed SNPs and can exploit LD between those markers and causal variants for tagging and prediction (Yang et al., 2010; Yang et al., 2015). Their heritability models nevertheless assign an expected contribution 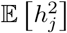 to each SNP and obtain total SNP heritability by summing these locus-level contributions (Speed, Holmes, and Balding, 2020). Such formulations omit the cross-locus terms *α*^*j*^*α*^*k*^*D*_*jk*_ that constitute the disequilibrium variance *C*_*t*_ (Campos, Sorensen, and Gianola, 2015; Lara et al., 2021). In populations under stabilizing selection, *C*_*t*_ is negative, so targeting only the genic variance and omitting disequilibrium will lead to an overestimate of genetic variance. At the posterior-median selection widths, the corresponding values are *C*_*t*_ */ V*_*t*_ ≈ −0.043 in African ancestry and 0.029 in European ancestry, or reductions of approximately 4.3% and 2.9%, respectively, in genetic variance relative to genic variance.

Reporting heritability should specify the population, time, and whether the target is the realized 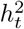 or an expectation across evolutionary histories. A pedigree-based mixed model is a good example: it assumes linkage equilibrium (no LD) and neutral evolution, and its target quantity resides in the founder population (*t =* 0) (Lynch and Walsh, 1998). The last condition keeps target parameters including genetic variance and heritability fixed, resolving the “realized– versus–expected” ambiguity. The same principle applies when attributing population differences in heritability or frequency-dependence to selection: doing so requires specifying the full evolutionary configuration (Coop and Przeworski, 2022; Bird, Jackson, and Winston, 2024; Schraiber and Edge, 2024; Huang et al., 2025). Nevertheless, current genomic approaches are largely implicit about these conditions.

### A generative model of frequency-dependent architecture

The SDE framework provides a practical way to generate allele frequencies and effects jointly under selection, recombination, drift, and demography. This improves upon benchmarks that simulate haplotypes under neutral demographic models and then assign effects using a phenomenological frequency-dependence model (Ashraf and Lawson, 2021; Taylor and Lawson, 2024). Assigning effects after simulating frequencies does not reproduce the feedback by which those effects influence allele survival under selection, including selection-induced LD. It therefore cannot by itself establish that the assigned architecture arose under the proposed evolutionary history.

Our approach jointly simulates the genetic background and the locus trajectories, fully capturing the evolutionary dynamics. We combined this simulation scheme with B-splines to capture the frequency-dependent architecture. This separates the evolutionary assumptions from the statistical procedure estimating the genetic architecture and provides a basis for simulation-based inference (Cranmer, Brehmer, and Louppe, 2020; Kim, Durvasula, and Zhang, 2025; Min et al., 2026). This generative approach overcomes known pathological behaviors of phenomenological models: *α*-models predict that squared effect size diverges as allele frequency approaches zero. This forced earlier studies to apply frequency thresholds in their analyses (Schoech et al., 2019; Rossen et al., 2026). To develop statistical methods, the next step is to combine this evolutionary generator with an explicit observation model. When the observation is a linear mixed model, one may follow the evolutionary mixed model presented in Lee and Terhorst (2026).

### Future directions and limitations

Our analytical results rely on an additive, highly polygenic trait and stabilizing selection that is broad relative to the additive genetic variance, *V*_*S*_ ≫ *V*_*t*_ + *C*_*t*_. Across the accepted posterior samples in the ABC analysis, the ratio (*V*_*t*_ + *C*_*t*_ ) *V*_*S*_ had present-day means of 0.0265 in African and 0.0176 in European ancestries, respectively; the corresponding 95th percentiles were 0.0287 and 0.0209. Thus, the approximation is accurate to within a few percent. An independent estimate also points to a large selection width relative to genetic variance, with an upper bound of (*V*_*t*_ + *C*_*t*_ ) / *V*_*S*_ ⩽ 1/28 ≈ 3.6% (Sanjak et al., 2017). The diffusion approximation for *C*_*t*_ does not enforce the biological constraint *V*_*t*_ + *C*_*t*_ ⩾ 0, because the continuous-time approximation does not respect the range constraints on LD (Kang and Rosenberg, 2019). We did not find any instances of this violation in our simulations. Parameter regimes in which it assigns appreciable probability to negative additive variance fall outside the useful range of this approximation and would require a formulation that preserves this constraint.

The model also treats the measured trait as the direct target of stabilizing selection, whereas human traits may instead be associated with fitness through pleiotropic effects on other traits (Simons et al., 2018; Koch et al., 2024; Simons et al., 2025). Extending the analysis to a focal trait that is only partially coupled to the selected phenotype would weaken its frequency-dependence, so the fitted selection widths and the magnitude of population contrasts could change. In particular, the selection width for fitness should generally decrease. It would not remove the main qualitative result that linkage and demography affect selection through the genetic background. Appendix E sketches the mathematical framework following Lee and Terhorst (2026).

The statistical uncertainty reported in the African-European frequency-dependence analysis accounts for evolutionary variation and ancestry-specific frequency-ascertainment effects. This includes the variation in the random effects model but ignores LDSC’s finite-sample estimation error. By allowing this estimation error, the posterior widths are likely to increase and wider ranges of the selection parameters *V*_*S*_ will become compatible with the observed frequency-dependence. Fully accounting for this uncertainty will require modeling the likelihood of GWAS summary statistics (Vilhjálmsson et al., 2015).

Population structure not in isolation is another important limitation not addressed in this work. We treat each ancestry as isolated and therefore omit migration among subpopulations. Recent work has studied the dynamics under stabilizing selection in the infinite-island setting (Li, Hermisson, and Sachdeva, 2026). It assumes a deterministic equilibrium and accounts for linkage disequilibrium accordingly, using the machinery of Veller and Coop (2024) and Negm and Veller (2026). Combining Li, Hermisson, and Sachdeva (2026)’s machinery with stochastic LD and nonequilibrium dynamics is an important future direction. This amounts to developing a stochastic recursion for disequilibrium variance in the presence of migration. Since admixture can be treated as a one-time migration event, its interaction with stabilizing selection should follow.

Temporal data or replicated populations would provide a complementary test of the model’s predictions (Schlötterer et al., 2014; Buffalo and Coop, 2019; Buffalo and Coop, 2020). The model predicts changes in LD and genetic variance, as well as the resulting frequency-dependence over time. Directly observing these quantities can help quantify the various evolutionary factors separately, providing information that contemporary summaries cannot offer, as shown in this work.

Efforts to recover evolutionary parameters from genome-wide association studies should focus on identifying parameters, or combinations thereof, that can be estimated from data (Sella and Barton, 2019). Although jointly estimating evolutionary forces from an association study alone might be impossible, certain nuisance parameters such as demography can be inferred from other sources like demographic inference (Spence et al., 2018; Kamm et al., 2019; Terhorst, 2025; Dilber et al., 2026). Hence, our work encourages researchers to explore holistic approaches combining statistical genetics methods and evolutionary inference to complement each other (Schraiber, Edge, and Pennell, 2024; Simons et al., 2025; Christ et al., 2025; Zhu et al., 2026; Lee et al., 2026).

## 4. Materials and Methods

### 4.1 Stochastic model and stationary distributions of macroscopic variables

We consider a population of *N*_*e*_ diploid individuals with a linear genome of size *L*. Mutation occurs at rate *u* per base pair per generation. We consider a single chromosome for the moment, but generalization to multiple chromosomes is straightforward. The trait mean and genic variance at time *t* are denoted by *m*_*t*_ and *V*_*t*_, respectively. We further consider the linkage disequilibrium (LD) variance *C*_*t*_. Suppose that individuals are assigned fitness proportional to

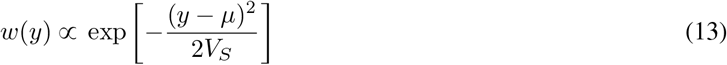

where *y* is the trait value. *V*_*S*_ > 0 is the selection width, which increases as selection weakens, and we assume that the phenotypic variance is much smaller than *V*_*S*_ (*V*_*S*_ ≫ *V*_*P*_ = *V*_*t*_+ *C*_*t*_+ *V*_*E*_) (Simons et al., 2018; Hayward and Sella, 2022; Simons et al., 2025). We also assume that the displacement *µ* is small relative to 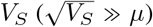. The environmental variance *V*_*E*_ is ignored because it can be absorbed into *V*_*S*_ (Simons et al., 2018; Negm and Veller, 2026). We assume that the trait is normally distributed. 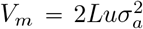 is the per-generation variance input from mutation, where 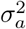 is the mutational variance. 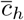 is the harmonic mean of the pairwise recombination fractions after replacing each *c*_*ij*_ with *c*_*ij*_ + 1/(2*N*_*e*_) − *c*_*ij*_/(2*N*_*e*_) to account for finite population size. We also require 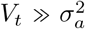, which follows from large *L* ≫ 1 and a small per-locus mutational variance 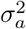.

Under these assumptions, we can write the stochastic differential equations (SDEs) of *m*_*t*_, *V*_*t*_, and *C*_*t*_ as

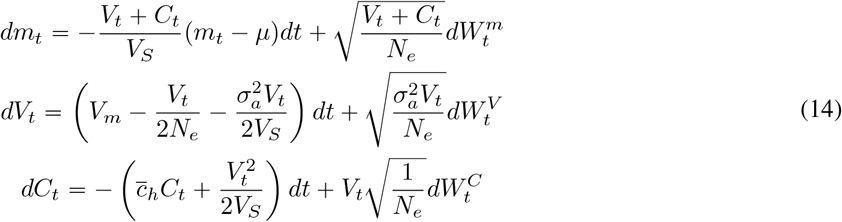

where 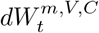 are independent Brownian motions.

Similar systems have been widely studied in the literature, and we justify each term of the SDEs in Appendix B (Barton, 1986; Barton and Turelli, 1987; Barton, 1989). The noise terms denoted by 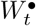 represent the random nature of these quantities. Biologically, the population mean is pulled toward the fitness optimum *µ* at a rate proportional to the additive genetic variance *V*_*t*_ + *C*_*t*_, while finite-population sampling adds random movement around this deterministic pull. Genic variance reflects a balance between new mutational input, loss by genetic drift, and loss by stabilizing selection. The LD component reflects recombination eroding existing disequilibrium, stabilizing selection continually generating negative disequilibrium, and drift introducing additional fluctuations. Therefore, the same parameters can still lead to different values of the trait mean and variance. Table 1 lists the aforementioned forces.

The SDE (14) generalizes the deterministic limits to stationary *distributions*. The genic variance process *V*_*t*_ has a Gamma stationary distribution with mean and variance

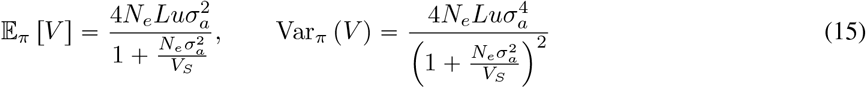

which coincides with the mutation-selection-drift balance limit under the stochastic House-of-Cards (SHoC) model (Bürger, Wagner, and Stettinger, 1989; Keightley and Hill, 1989; Bürger and Lande, 1994; Ragsdale, 2026). For the LD process, although a closed-form stationary distribution is not available, its mean is

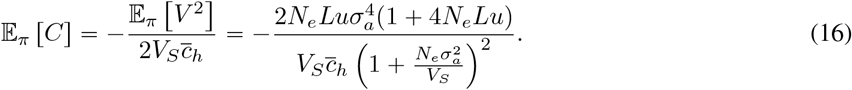

Finally, the mean process *m*_*t*_ has the stationary distribution

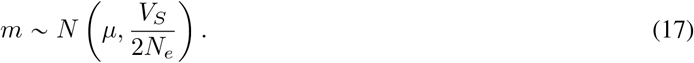

When the optimum is centered at zero, this reduces to the expression *N (*0, *V*_*S*_ */* 2*N*_*e*_)) used in the simulations. The derivation of the stationary moments is in Appendix C.

An important mathematical device appearing in Negm and Veller (2026) is the proportionality constant *K* that relates genic variance *V*_*t*_ and disequilibrium *C*_*t*_. They explained how either *K* or *C*_*t*_ can be computed from the other, but did not show how to obtain either one from the evolutionary parameters of the system. We show how this can be done in Appendix B by jointly modeling *V*_*t*_’s time evolution alongside *C*_*t*_, making the genetic system self-contained. This development complements Negm and Veller (2026)’s formula, which predicts allele-frequency dynamics conditional on the trait’s genetic variance: our SDE provides the temporal trajectory of *C*_*t*_ and *V*_*t*_ on which those dynamics depend. This lets us simulate the system from scratch and under changing demography, allowing heritability and frequency-dependence architecture to emerge as outputs of the model rather than an assumed input.

### 4.2 Deterministic and averaged locus dynamics

To allow the genetic background of a focal allele to change over time, we write the dynamics of *x*^*j*^ as

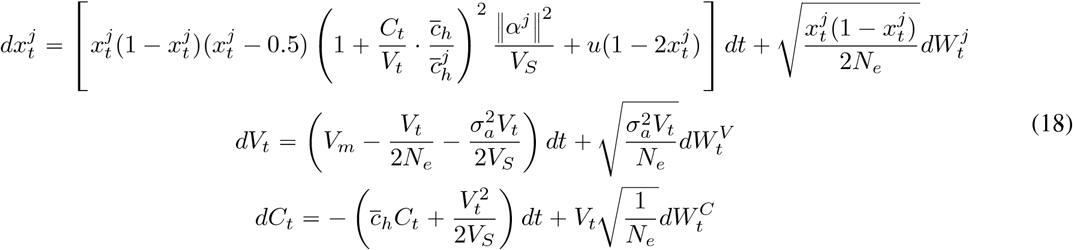

We assume that 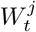 is independent of 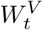 and 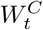. Here, 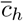 and 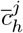 use the finite-population-adjusted pairwise fractions *c*_*jk*_ + 1 / (2*N*_*e*_) − *c*_*jk*_/ ( 2*N*_*e*_ ). The first equation describes how the frequency of a focal allele responds to stabilizing selection in the genetic background summarized by *V*_*t*_ and *C*_*t*_. The selection term depends on the allele’s current frequency, its effect size, and the LD correction *C*_*t*_ */ V*_*t*_, while symmetric mutation contributes the term 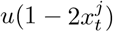 and genetic drift contributes the Brownian noise. The latter two equations are the same *V*_*t*_ and *C*_*t*_ components of the macroscopic SDE in equation (14), repeated here to emphasize that the focal allele is coupled to the evolving population-level variance and disequilibrium. Treating 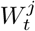 as independent of 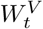 and 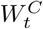 is a plausible approximation because the contribution of a single locus to the macroscopic variables *V*_*t*_ and *C*_*t*_ is negligible.

For comparison with the stochastic finite-population model, we define a deterministic trajectory using the average population moments E _*π*_ [*V*_*t*_] and E_*π*_ [*C*_*t*_] of the full stochastic system,

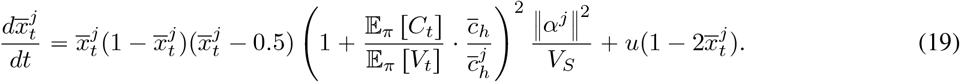

This substitutes *V*_*t*_ and *C*_*t*_ with their averages under the stationary law *π*. For the deterministic denominator of equation (8), these stationary moments are computed from the recursion using the raw harmonic mean in the infinite-population limit studied by Veller and Coop (2024) and Negm and Veller (2026). They therefore differ from the stationary moments in the stochastic numerator, whose recursion uses the finite-population-adjusted harmonic mean. If this trajectory is viewed as the drift component of 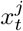’s SDE, the correct drift coefficient capturing the effective strength of selection under the polygenic assumption is

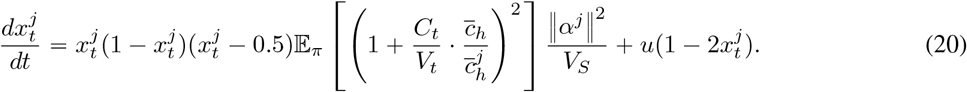

The proof is based on the polygenic assumption and is provided in Appendix D.

### 4.3 SLiM simulation configuration

We used three SLiM-based individual-level simulation workflows to evaluate the SDEs. The constant-demography workflows model a diploid population of size *N*_*e*_ *=* 1000 evolving along a genome of length *L =* 10^8^. For the locus- and background-level validation under stabilizing selection, we used the per-base mutation rate *u =* 10^−10^ and drew mutational effects from a normal distribution with variance 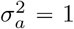. For each run, we recorded the population mean, the total genetic variance, and the genic variance at every generation. Genic variance was computed by summing 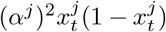, and genetic variance was obtained from the variance of genetic values. Disequilibrium was taken as the difference between the two quantities.

Figure 1 was used to isolate the contribution of disequilibrium to short-term fluctuations in additive variance under neutrality. These simulations used *N*_*e*_ *=* 500, a uniform per-base recombination rate of *r =* 10^−8^, and were run for 2,000 generations with 100 independent replicates. Figure A1 was used to compare the proposed SDE system with individual-based simulations under Gaussian stabilizing selection. Here, fitness was centered at the optimum *µ* = 0 with selection width *V*_*S*_, and we considered the grid *V*_*S*_ ∈ {500, 1000, 2000} together with recombination rates *r* ∈ {10^−8^, 10^−7^, 10^−6^}. Each parameter combination was simulated for 8,000 generations with 100 independent replicates. These runs were used to compare the terminal distributions of *m*_*t*_, *V*_*t*_, and *C*_*t*_ inferred from SLiM, where *C*_*t*_ was obtained as the difference between total genetic variance and genic variance, against the corresponding distributions generated by the SDE approximation.

We repeated the stabilizing-selection comparison under a modified version of the `Zigzag_1S14` demography of Schiffels and Durbin (2014). To retain its three expansions and two declines, we rescaled its oldest time point to generation 8,000 and preserved the original factor-of-four spacing between backward-time breakpoints. Population size alternated between *N*_*e*_ *=* 500 and *N*_*e*_ *=* 2,000, changing exponentially within each interval, and the same generation-specific integer population sizes were supplied to SLiM and the SDE. We used 100 replicates for each combination of *V*_*S*_ ∈{1000, 2000, 4000} and *r* ∈ {10^−8^, 10^−7^, 10^−6^}.

### 4.4 Frequency-dependence analysis with B-splines

In these experiments, we set 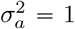 and *u* = 10^−10^ to increase the selection experienced by loci. This is equivalent to *u* = 10^−8^ with polygenicity of 1%. To estimate the frequency-dependent architecture from the SDE simulations, we modeled the conditional mean of squared effect size, E [(*α*^*j*^) ^2^ | *x*^*j*^ (1 − *x*^*j*^)], using the simulated pairs (*α*^*j*^, *x*^*j*^) recorded at the final generation. For each parameter configuration under constant demography, we used 4 × 10^6^ replicates and treated heterozygosity *x*^*j*^ (1 − *x*^*j*^) as the predictor and (*α*^*j*^ )^2^ as the response. The regression function was represented by a degree 5 B-spline with 20 interior knots placed uniformly over the observed range of heterozygosity. Spline predictions were then evaluated on a grid of 200 equally spaced heterozygosity values. To preserve the expected decline in effect size with allele frequency, we imposed shape restrictions so that the fitted spline is non-increasing and convex (Ramsay, 1988; Meyer, 2008). Operationally, these constraints were enforced through a reparameterization of the spline coefficients and fitting by nonnegative linear least squares using `scipy`.

For the non-equilibrium demography, we used the human demographic model `OutOfAfrica_3G09` of Gutenkunst et al. (2009) as implemented in `stdpopsim` (Lauterbur et al., 2023). For each present-day subpopulation, this demographic model was converted into a sequence of lineage-specific time intervals with piecewise changes in effective population size discretized at the generation level. Equation (18) was then simulated generation by generation along each such trajectory, so that the deterministic and stochastic terms evolved with the time-varying *N*_*e*_. The harmonic mean recombination fraction was also updated by generation: for each contemporaneous *N*_*e*_, pairwise recombination fractions were replaced with *c*_*ij*_ + 1 / (2*N*_*e*_) − *c*_*ij*_ */* 2*N*_*e*_ before averaging over the expected number of segregating loci. For computational efficiency, these harmonic means were evaluated on a log-spaced grid of population sizes and interpolated along each generation-level demographic trajectory. Initial allele frequencies were sampled from the neutral beta distribution implied by the deepest ancestral effective population size, while *V* was initialized from its stationary distribution and *C* from its stationary mean under that ancestral state. After these simulations, the same constrained spline-fitting procedure was applied separately within each subpopulation, yielding demographic-specific estimates of E [(*α*^*j*^) ^2^ | *x*^*j*^ (1 − *x*^*j*^)] without assuming stationarity.

### 4.5 Approximate Bayesian computation on GWAS summary statistics

To compare the predicted population differences with empirical estimates, we extracted the meta-analysed conditional squared effect sizes and corresponding minor allele frequencies reported by Rossen et al. (2026). These estimates were obtained with stratified LD score regression (S-LDSC), which partitions per-SNP heritability according to allele frequency (Finucane et al., 2015). For each allele-frequency bin, the corresponding S-LDSC coefficient estimates the average per-SNP heritability in that bin, which converts to normalized average squared effect size (Rossen et al., 2026). These conditional squared-effect-size averages were used as the summary statistics for the approximate Bayesian computation (ABC).

The variants in the African frequency bins are conditional on being common (MAF ⩾ 0.05) in the European ancestry, and vice versa. Rossen et al. (2026) reported five quintile frequency bins in both ancestries. For each bin, we fitted

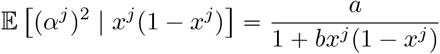

where *x*^*j*^ is the minor allele frequency. Taking reciprocals gives

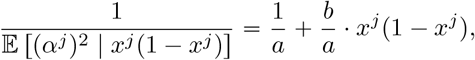

so we estimated the intercept 1 / *a* and the slope *b* / *a* with ordinary least squares. The target quantity *b* was obtained by taking the quotient of the slope and the intercept. The normalization applied to the squared effect sizes only changes *a*, so the quotient *b* = (*b*/*a*)/(1/*a*) is unaffected. These steps were applied to both ancestries, resulting in *b*_AFR_ and *b*_EUR_.

ABC requires modeling the data-generating process that precedes the summary-statistic calculation described in the previous paragraphs. As noted in the main text, E [(*α*^*j*^) ^2^ | *x*^*j*^ (1 − *x*^*j*^)] emerges from the evolutionary process. To account for the demographic effect alongside the evolving background, we extend Patel et al. (2024)’s conditional frequency spectra approach. We replaced the fastDTWF (Spence et al., 2023) engine with the diffusion approximation and further accounted for frequency ascertainment using the Kolmogorov equations (Pavliotis, 2014).

Using `OutOfAfrica_3G09` demography without migration, we sampled the history ℋ = {*V*_*t*_, *C*_*t*_, *N*_*e*_ *(t)*} from the ancestral population down to the present African and European populations. The distribution of *x*^*j*^ conditional on *α*^*j*^ and is determined by the locus SDE (5). The shared ancestry between African and European populations is accounted for by the distribution of *x*^*j*^ at the time of the population split *t*_*s*_. This distribution is obtained by solving the forward Kolmogorov equation corresponding to the SDE (5). The distribution of allele frequency ascertainment conditional on *x*^*j*^ at *t*_*s*_ is then obtained by solving the backward Kolmogorov equations starting at present time and moving back up to the split time *t*_*s*_. We can then use Bayes’ rule by integrating over these probabilities to obtain the squared effect size (*α)*^*j* 2^ of each frequency bin conditional on the history ℋ. The mathematical details are provided in Appendix F.

We inferred two selection widths: *V*_*S*,AFR_ for both the shared ancestral history and the African descendant history, and *V*_*S*,EUR_ for the European descendant history after the split (Beaumont, Zhang, and Balding, 2002). Thus, *V*_*S*,anc_ *V*_*S*,AFR_. Here, a particle refers to a draw of the parameter pair *V*_*S*,AFR_, *V*_*S*,EUR_ and the subsequent simulation data generated from it, including the history ℋ and the two fitted coefficients *b*_AFR_ and *b*_EUR_. We drew the two selection widths independently from continuous uniform priors on [2000, 8000] .

Acceptance of a particle was based on the covariance-weighted Euclidean distance between simulation and Rossen-fitted **b** = (*b*_AFR_, *b*_EUR_) estimates. That is, for a covariance matrix Σ estimated from a draw of pilot particles, the distance was defined as the square root of

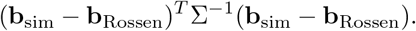

A total of 10,000 pilot particles were used, and we added a 10^−8^ tr (Σ) / 2 times the identity matrix to Σ to keep it invertible. The rejection tolerance was the 0.5th percentile of the pilot joint distances. We then applied this fixed tolerance to 100,000 independently simulated production particles and accepted 374 particles (0.374%).

For the posterior-predictive curves in Figure 4A, we sampled 500 accepted joint particles with replacement. Within each population, we multiplied every predicted bin in a draw by a single constant to align the *y*-intercept. Recall that this scaling does not affect *b* estimates. This adjustment calibrates only the vertical scale *a* and leaves the simulated frequency-dependence coefficient *b* unchanged. Pointwise medians and the 2.5% and 97.5% quantiles across the 500 draws give the displayed posterior-predictive curves and intervals. They incorporate posterior uncertainty in both selection widths and variation among shared evolutionary histories, but not uncertainty from GWAS phenotype sampling and LDSC estimation error.

### 4.6 Human linkage map

We used the HapMap Phase II GRCh37 genetic map implemented in `stdpopsim`. For each focal locus *j*, the finite-population-adjusted local harmonic mean of equation (4) is defined over the *L* − 1 loci pairs as

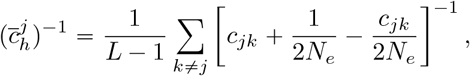

where *c*_*jk*_ = [1 − exp(−2*d*_*jk*_)]/2 is Haldane’s map function of the HapMap genetic distance *d*_*jk*_. The genome-wide harmonic mean of equation (4) is the harmonic mean of the locus-specific values, 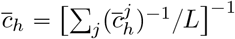; we estimated the outer average from 3,000 focal loci sampled across the autosomes in proportion to physical interval length, for each *N*_*e*_ ∈ {1000, 7300, 50000}.

## Code availability

The code implementing the experiments presented in the paper is available at https://github.com/hanbin973/hlee_sto_bulmer.

## Acknowledgments

We thank Joshua Schraiber for discussions on the Bulmer effect. This research was supported in part by the National Institute of General Medical Sciences of the NIH under award number R35GM151145. The content is solely the responsibility of the authors and does not necessarily represent the official views of the NIH.

## A Historical note

By assuming that the variance remains constant at the equilibrium value, one can show that the trait mean follows the Ornstein-Uhlenbeck (OU) process with an attracting force toward the fitness optimum (Lande, 1976). This approximation is effective because variance changes more slowly than the mean. Therefore, over a suitably short period, the change in the mean can be analyzed while holding the variance fixed. This formulation leads to the classic Gaussian stationary distribution of trait mean, generalizing the deterministic equilibrium to the stationary distribution.

Attempts to further generalize to random trait variance have mostly taken bottom-up approaches, where macroscopic quantities such as the trait mean are deduced from microscopic states of segregating alleles (Bürger, Wagner, and Stettinger, 1989; Barton, 1989; Walsh and Lynch, 2018; Bod’ová, Tkačik, and Barton, 2016; Bod’ová, Szép, and Barton, 2021). One starts from the stationary distribution of alleles and takes the product of these distributions under the *linkage-equilibrium* (LE) assumption – that is, correlations between variants are absent. The trait’s stationary distribution then follows from the joint distribution of microscopic states. In general, the precise distribution of the trait variance has no closed-form solution, so only the first few moments are available through series-expansion techniques. Furthermore, the LE assumption ignores the genetic variance reduction by negative linkage disequilibrium (LD) due to stabilizing selection, known as the Bulmer effect.

A natural relaxation of the LE assumption is to first obtain the deterministic steady-state LD and plug it into the microscopic (locus-level) dynamics to obtain a refined stationary distribution. Then we obtain the stationary distribution of macroscopic states as a function of the microscopic variables. In general, even in the deterministic case, the dynamics of population LD are extremely difficult to solve (Burger, 2000). Fortunately, quasi-linkage equilibrium (QLE) is a moment-closure technique that assumes weak selection relative to recombination and greatly simplifies the system (Kimura, 1965). Lande (1975) derived the formula for steady-state LD under a Gaussian allelic-effect assumption, and Barton and Turelli (1991) further generalized it to non-Gaussian settings. The most general treatment in the multilocus setting was presented by Nagylaki (1993). However, the application of these steady-state LD formulas to locus-level dynamics was only recently developed by Negm and Veller (2026). Lee and Terhorst (2026) used a heuristic technique based on the results of Negm and Veller (2026), but obtained results only for the population’s *genic* variance and not the entire *genetic* variance, thereby ignoring LD’s contribution to genetic variance. In fact, this result for genic variance coincides with the earlier work of Keightley and Hill (1989) and Barton (1989), who assumed LE (see also Bürger and Lande, 1994).

## B Deriving the stochastic differential equations

The results are derived from a discrete-time model and then approximated by a diffusion using the infinitesimal mean and variance. The derivations are largely heuristic and were validated primarily by individual-based simulations rather than by rigorous mathematical proofs. The mean SDE is standard, so we refer readers to Lande (1976). The genic variance SDE was introduced by Barton (1986), Barton and Turelli (1987), and Barton (1989), so we omit the derivation here. We deduce the SDE for disequilibrium *C*_*t*_ in this appendix.

Let *C*_*t*_ be the linkage disequilibrium (LD) component of additive variance

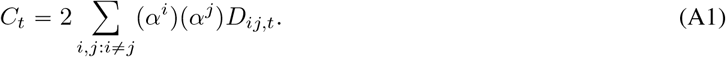

Each component *D*_*ij,t*_ for the locus pair (*i, j*) is discounted by a factor of *c*_*ij*_ every generation

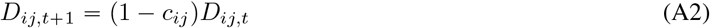

in an infinite population with no selection, ignoring the drift noise. Note that we assume random mating that eliminates trans-LD at each generation after mating. Therefore, we only keep track of cis-LD as *D*_*ij,t*_. We then follow the life cycle

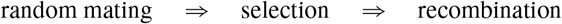

life cycle. If selection creates total LD of 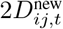 at each generation, this LD is equally divided between cis- and trans-LD. Therefore, the recursion is

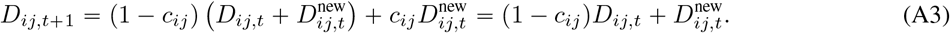

Note that an extra 1 − 1/(2*N*_*e*_) factor contributes to the decay in finite populations

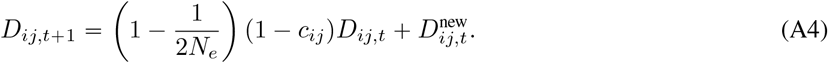

Bulmer (1971) showed that the total amount of new (cis-)LD generated each generation is

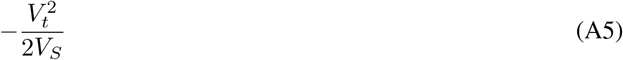

under stabilizing selection with selection width *V*_*S*_. In Bulmer (1974), Bulmer assumed a varying recombination fraction *c*_*ij*_ but a uniform effect size *α*^*i*^ *α* across all loci. As a result, each (*i, j)* receives an equal share of the total LD, giving

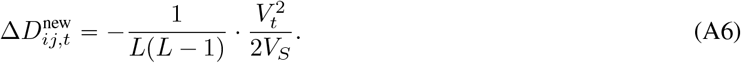

Note that Bulmer (1974) used *L( L* − 1 )/2 unordered pairs *i* < *j* whereas this work considers *L* (*L* − 1) ordered pairs (*i, j*) . This choice is entirely a matter of notation and has no material consequences.

Lande (1975) showed that when the *α*^*i*^ values are also allowed to vary, each pair (*i, j*) divides the total new LD in proportion to

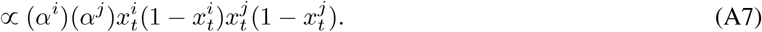

This follows from the fact that the trait disequilibrium (*α*^*i*^)(*α*^*j*^)*D*_*ij,t*_ is proportional to

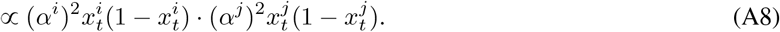

This aligns with our intuition: locus pairs with larger effect sizes (*α*^*i*^)(*α*^*j*^) contribute more to selection-induced LD. It also suggests that pairs with larger variance 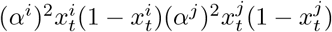 contribute more to LD.

Assuming the polygenic model in which a single locus occupies a negligible amount of genic variance compared to the aggregate, equation (A6) generalizes to

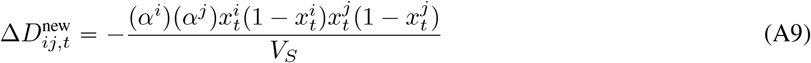

which can be verified by

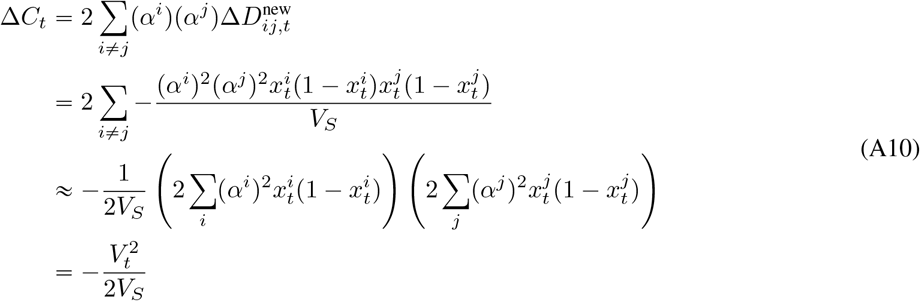

and

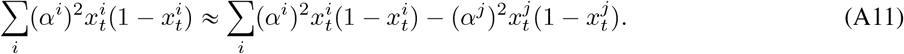

Substituting this formula into equation (A4) gives

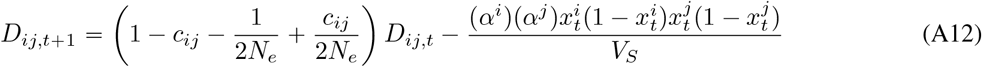

which has a steady-state solution (Δ*D*_*ij,t*_ = 0)

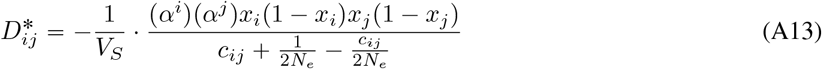

which coincides with that of Negm and Veller (2026) when *N*_*e*_ → ∞ (we show the equivalence shortly). This is a quasi-steady-state statement, as *x*_*i*_ *x*^*i*^ is still changing over time. In fact, stabilizing selection does not permit a stable equilibrium for *x*_*i*_. A more precise statement is that LD dynamics are faster than allele-frequency dynamics, so LD can be treated as a dependent variable of the latter (Berglund and Gentz, 2010).

Negm and Veller (2026) cite Lande (1975) to write

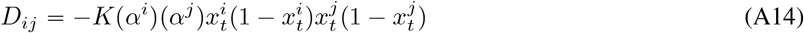

for some constant *K* and found that

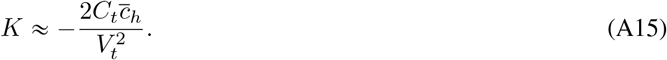

With 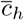 as the harmonic mean of recombination fractions, equation (A13) suggests that

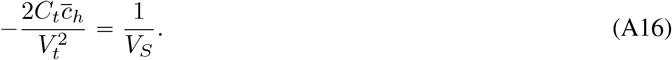

This can be shown by substituting the quasi-steady-state disequilibrium

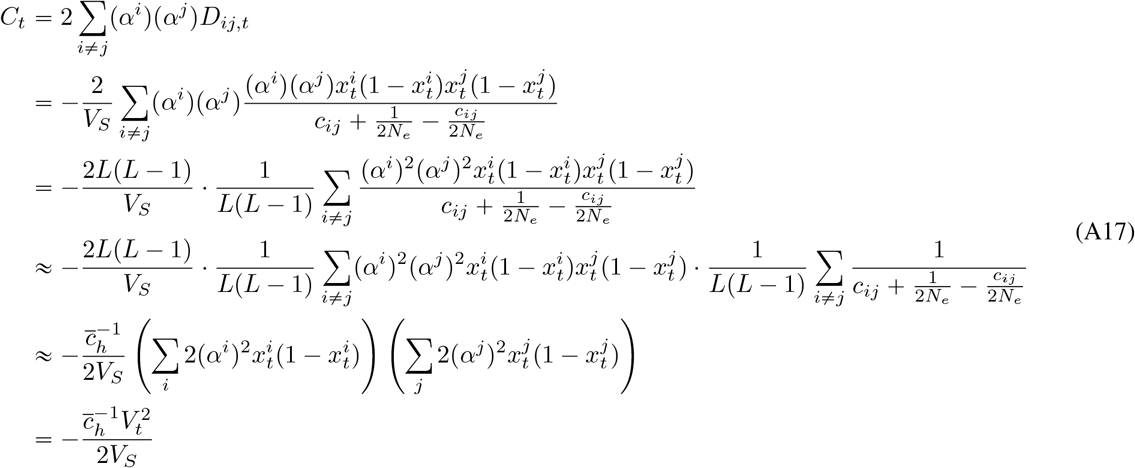

with

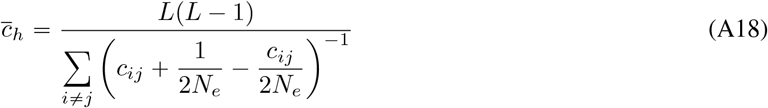

being the harmonic mean of the finite-population-adjusted recombination fractions. In particular,

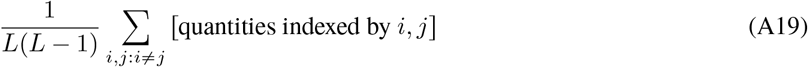

was treated as an average over *L*(*L* − 1) pairs, invoking E [*XY* ] = E [*X*] E [*Y* ].

In a similar fashion, we can compute the disequilibrium *C*_*t*+1_ in the next generation. Summing over *D*_*ij,t*±1_ gives

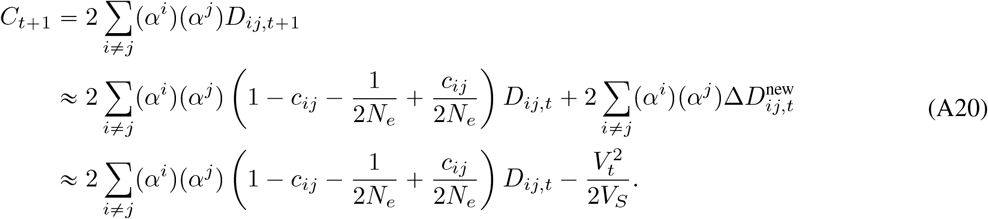

We further expand the first sum using a quasi-steady-state approximation (without the 2 factor)

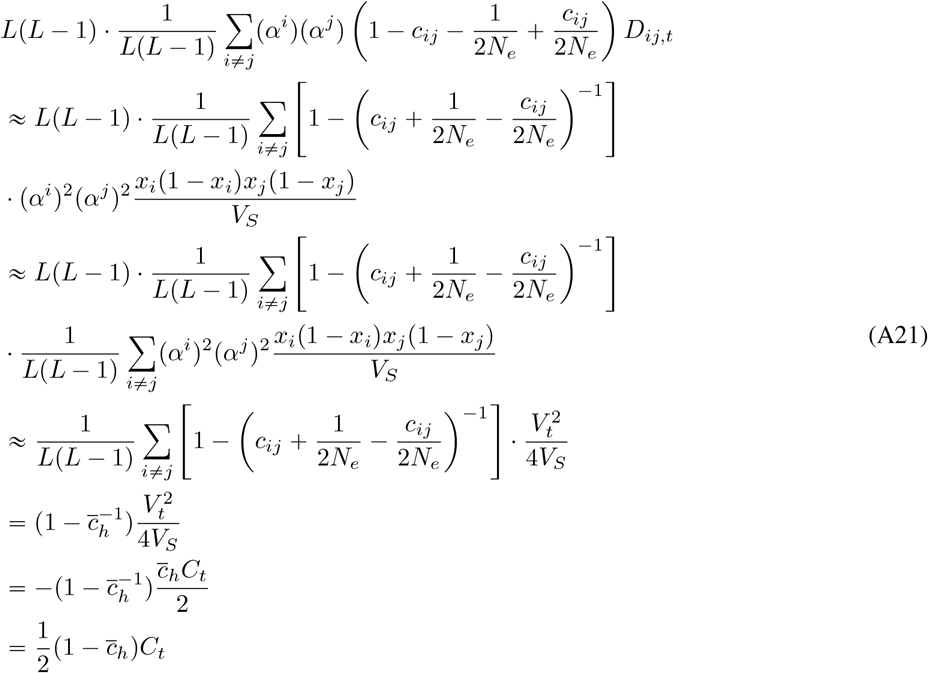

Putting this back into *C*_*t*±1_ gives

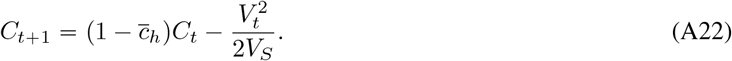

To account for finite-population noise, Ohta and Kimura (1971) obtained the single-generation variance in LD for loci *i* and *j* as

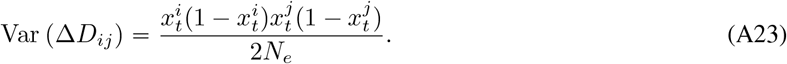

We substitute this expression into

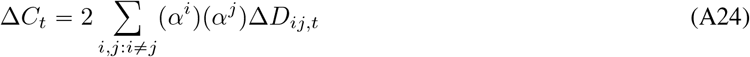

and obtain

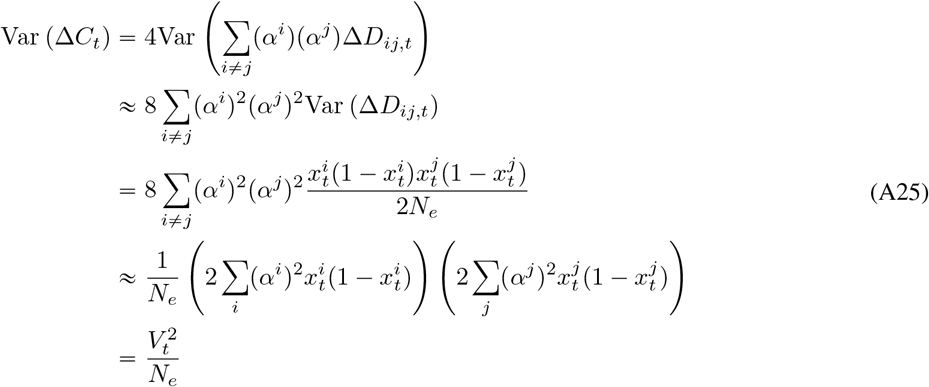

where we assumed in the second line that Cov (Δ*D*_*ij,t*_, Δ*D*_*kl,t*_) = 0 for all quadruplets (*i, j, k, l*) and in the fourth line that 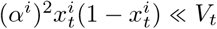.

Combining the results so far, we have

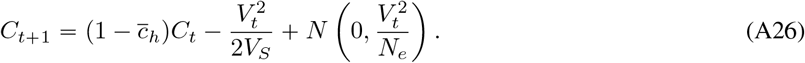

This has an expected steady-state value of

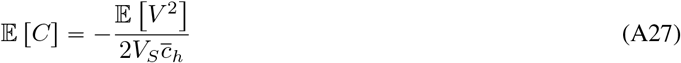

by taking expectations on both sides and assuming the existence of a stationary distribution. This is approximately the same as equation (A17) for large *N*_*e*_. The continuous-time approximation is

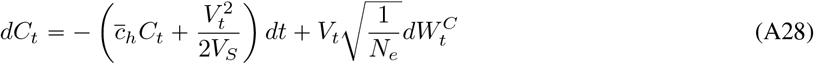

which is identical to the equation in the main text.

## C The stationary moments

We give the derivation of the stationary quantities used in the main text. The genic variance process has the form

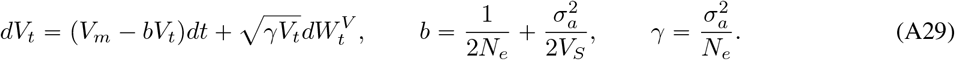

This is a Cox-Ingersoll-Ross (CIR) diffusion on R_⩾0_. Its stationary density *π*_*V*_ is obtained by solving the one-dimensional stationary Fokker-Planck equation under zero probability flux,

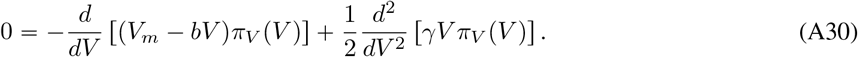

The unique solution is the Gamma density

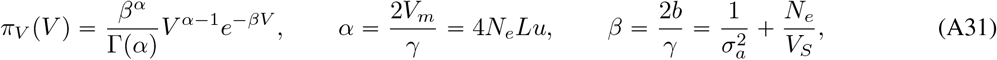

for *V* > 0, whose mean and variance are given in equation (15).

For the LD process, the stationary mean follows by taking expectations in

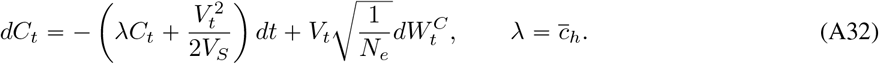

Assuming the stationary distribution exists,

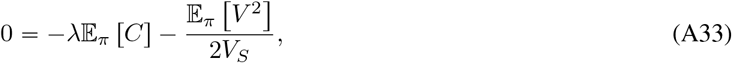

which gives equation (16). Higher-order moments of *C*_*t*_ are also available in closed form, but are not used here.

The dynamics of the expected variances follow directly from the SDEs. Taking expectations in the *V*_*t*_ and *C*_*t*_ equations and using the fact that the stochastic integral terms involving 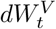 and 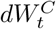 have mean zero gives

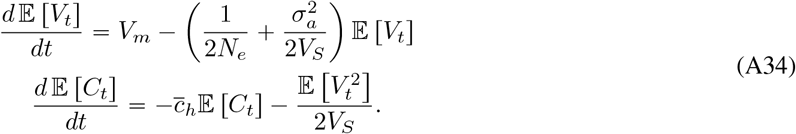

We next justify why random fluctuations in *V*_*t*_ and *C*_*t*_ do not change the stationary distribution of the trait mean.

From equation (14),

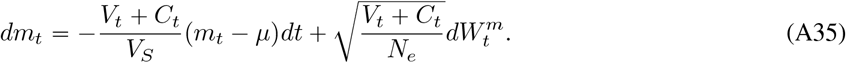

Writing

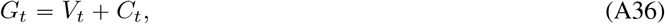

the mean equation is

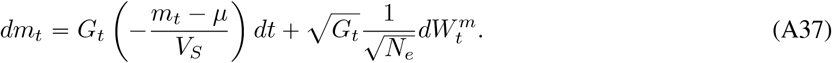

As long as the additive variance *G*_*t*_ is positive, this is an OU process run on the random clock (Øksendal, 2003)

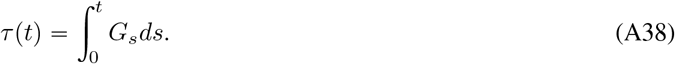

Equivalently, the generator acting on smooth test functions *f* (*m*) is

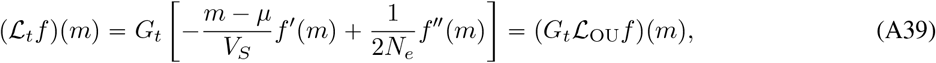

where ℒ_OU_ is the generator of the OU process

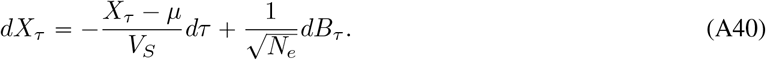

The factor *G*_*t*_ *= V*_*t*_ + *C*_*t*_ changes only the speed at which the process moves along this OU trajectory. It therefore preserves the eigenfunctions of _OU_ and rescales the corresponding eigenvalues by a positive factor. In particular, the stationary density *π*_*m*_ solves

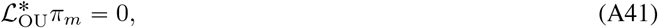

and hence also solves 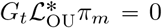, where the asterisk denotes the adjoint operator. Consequently, the stationary law of the mean is the OU stationary law

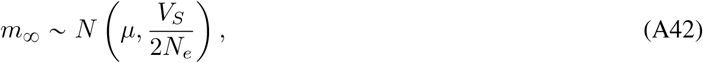

independent of the stationary fluctuations of *V*_*t*_ and *C*_*t*_.

## D The average locus dynamics

We prove the averaged dynamics by showing that the density of *x*^*j*^ approximately satisfies the following Fokker-Planck equation (FPE)

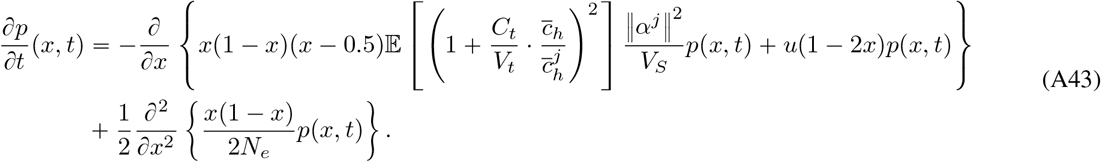

In doing so, we introduce the generator for Markov processes (Pavliotis, 2014). The adjoint generator 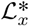 of the Markov process associated with the above equation is

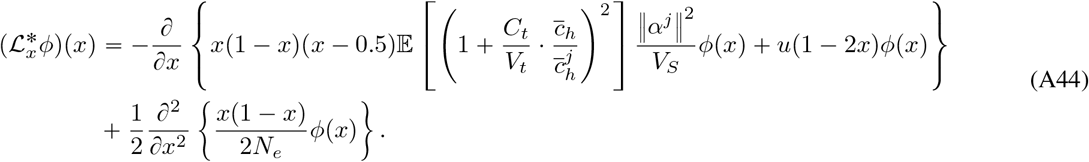

acting on smooth probability densities *ϕ x* . Then, the FPE can be written as 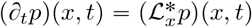. We show that for any smooth function *ψ*, the FPE for *x*^*j*^ holds in the distributional sense

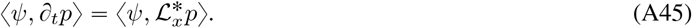

with the inner product defined by integration

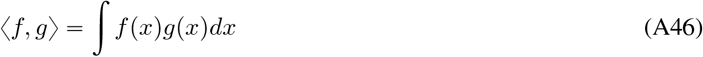

and the density *p*(*x, t*) of 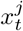.

Let ℒ ^∗^ be the adjoint generator of the full process (*x*^*j*^, *V, C*) in equation (18).

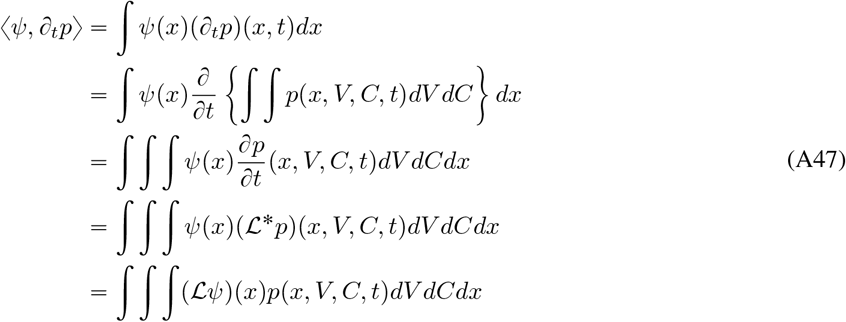

where in the last line we used the adjoint property

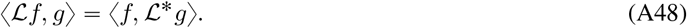

From the definition of the generator ℒ of equation (18), we have

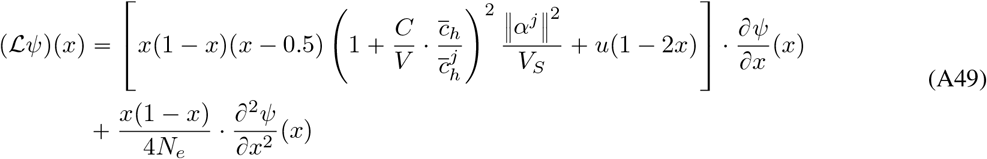

because the derivatives with respect to *V* and *C* all vanish, as *ψ* depends only on *x*. The inner integration with respect to *V* and *C* in equation (A47) is

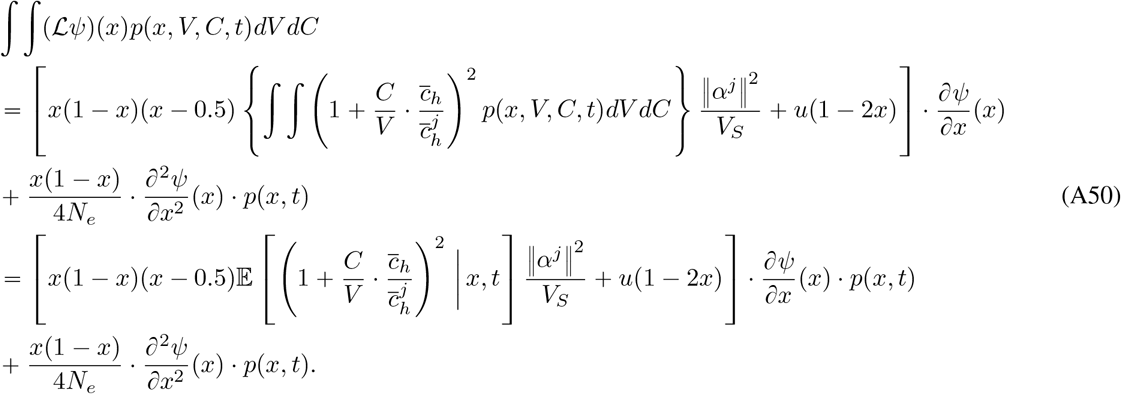

The most important step is to invoke the polygenic assumption to claim

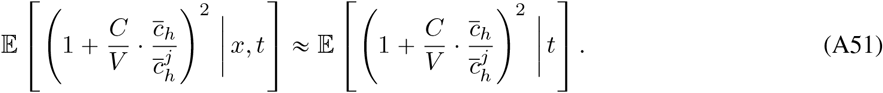

This approximation says that the distribution of *V* and *C* depends very little on *x* because an individual locus contributes nearly nothing to the macroscopic variables.

Dropping the conditional mean’s dependence on *x* simplifies equation (A47) to

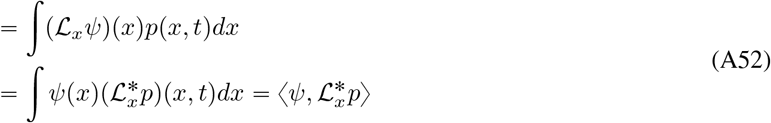

yielding the desired result.

## E Pleiotropic effects on a focal trait under changing demography

Following Lee and Terhorst (2026), we separate the evolutionary distribution of effects on traits under selection from their influence on a focal observed trait. Let *α* be the vector of allelic effects across traits under stabilizing selection, and let *β* be the effect of the same allele on the focal trait. The squared magnitude ‖*α*‖ ^2^, for the single selected trait in the main text, reduces to *α*^2^. Write allele frequency as *x*, suppressing the time index throughout this section. The distribution being conditioned on may correspond to any generation and need not be stationary.

The conditional second-moment model of Lee and Terhorst (2026) is

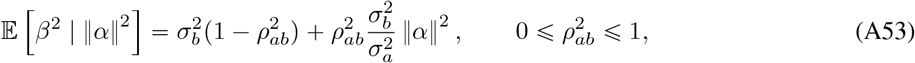

where 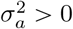 is the expectation of 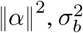 is the second moment of the focal-trait effect, and 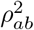 controls the coupling between the two.

This formulation includes a focal trait that is one of several traits under selection, as in multivariate models of stabilizing selection (Simons et al., 2018; Simons et al., 2025). For example, if *α* has *d* exchangeable components and *β* is one of them, symmetry gives

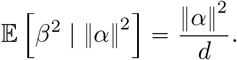

As explained by Lee and Terhorst (2026) for the isotropic model of Simons et al. (2018), this is recovered by setting 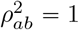 and 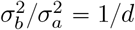.

Assume that the focal-trait effect provides no additional information about allele frequency once the squared magnitude of the selected effect is specified:

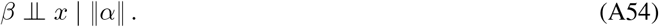

This can be justified by the graphical model

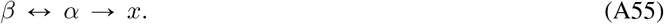

Evolution reweights the distribution of selected-effect magnitudes. The law of total expectation then gives

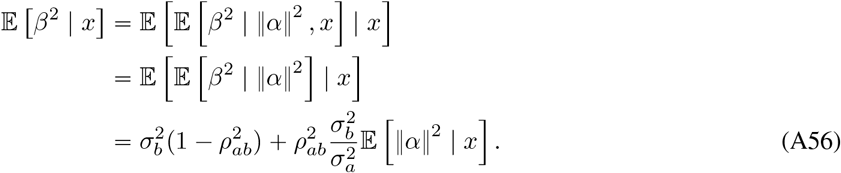

Therefore, the focal trait’s frequency-dependence is a linear function of E [‖*α*‖^2^ | *x*] . The calculation requires neither stationarity nor linkage equilibrium. Demographic history and stochastic linkage disequilibrium enter through the evolutionary conditional moment E [‖*α*‖^2^ | *x*].

## F Forward–backward reconstruction of empirical frequency bins

We describe the calculation used to reconstruct the cross-population frequency bins reported by Rossen et al. (2026). Write 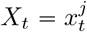 for the frequency of a focal allele and *a*^2^ for a possible value of its squared effect (*α*^*j*^)^2^. Let

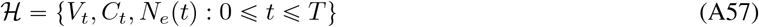

denote one realized genome-wide evolutionary history. The history contains a common ancestral path until the split between African and European ancestries at time *t*_*s*_ and two independently evolving descendant paths thereafter. Conditional on (*α*^*j*^)^2^ = *a*^2^ and ℋ, equation (18) has backward generator

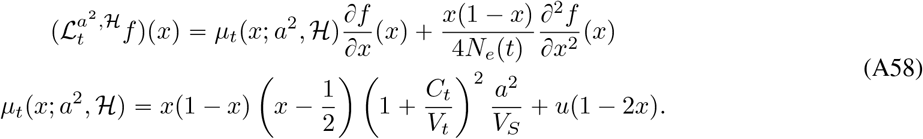

Here we use the simplifying assumption 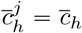. The focal effect is independent of the macroscopic history under the polygenic approximation, 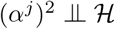.

The calculation is a history-conditioned version of the forward–backward decomposition used to construct conditional frequency spectra (Patel et al., 2024). Let *ρ*_*t*_ (*x* | *a*^2^, ℋ) be the probability density of *X*_*t*_ along the ancestral branch. Starting from the neutral beta distribution implied by the ancestral population size and mutation rate, we propagate this density forward to the split *t*_*s*_ using the Fokker–Planck (forward Kolmogorov) equation

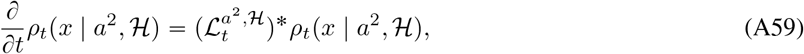

where 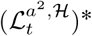 is the adjoint of the generator in equation (A58).

For a descendant population *P* P tAFR, EURu and minor-allele-frequency bin *B*_*k*_, define

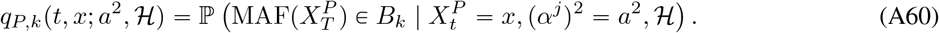

This probability is propagated backward along the trajectory for population *P* by

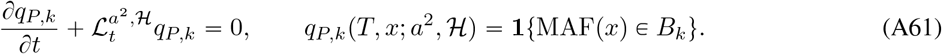

We similarly define *q*_*P,G*_ using the terminal event *G* = { *x* : MAF(*x*) > 0.0499} that a variant is common in population *P*.

At the population split, descendant allele-frequency paths are conditionally independent given their common ancestral frequency *X*_*t*_, the squared effect *a*^2^, and ℋ. Let *A*_AFR,*k*_ be the event that a variant lies in African-ancestry bin *B*_*k*_ and is common in European ancestry. Conditioning first on the ancestral frequency at the split gives

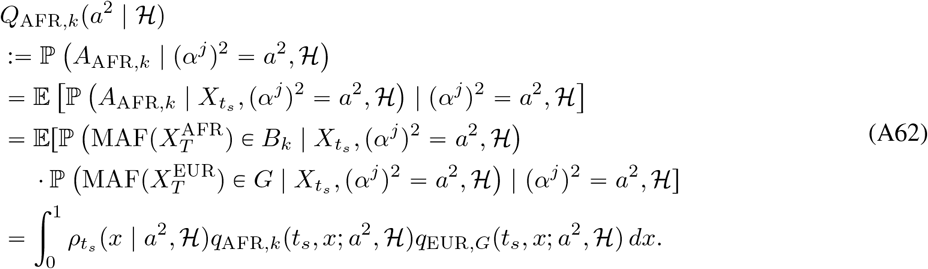

The European-ancestry-bin probability *Q*_EUR,*k*_ exchanges the roles of African and European ancestry in equation (A62). This product of backward probabilities is where the two present-day frequency filters combine while retaining their shared ancestral frequency.

Let *g (a*^2^) be the density of (*α*^*j*^) ^2^ induced by 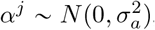, and let *A*_*P,k*_ denote the corresponding joint ascertainment event for population *P* . Because (*α*^*j*^) ^2^ ╨ ℋ, its conditional density given ℋ is also *g* (*a*^2^ ). The ascertainment probability conditional on the shared history is therefore

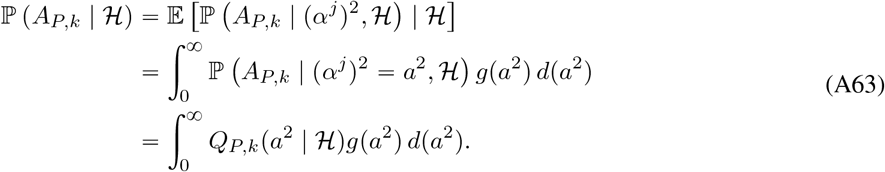

For *p* = 1, 2, the corresponding ascertainment-weighted joint moment follows from the same conditioning argument:

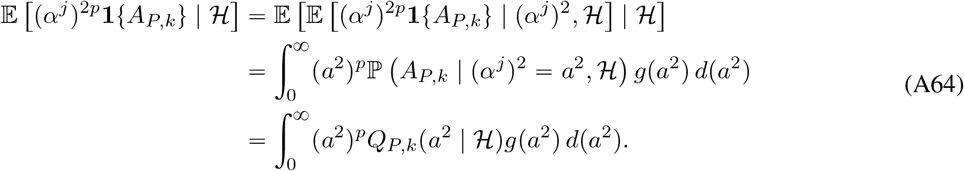

Dividing equation (A64) by equation (A63) gives the conditional moments needed for the bin mean and variance:

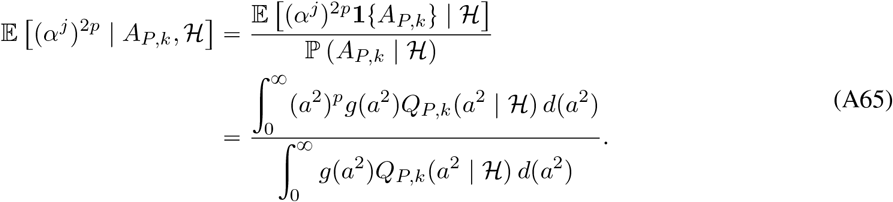

In particular, write

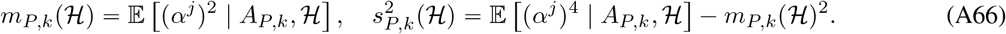

We evaluate the integrals in equation (A65) directly by generalized Gauss–Laguerre quadrature, avoiding brute-force simulation of enough loci to populate every joint frequency bin.

For a bin containing *n*_*P,k*_ independently sampled ascertained variants, let

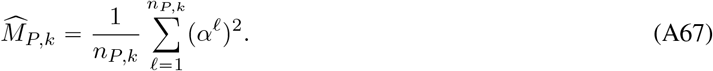

The law of total covariance separates variation among shared evolutionary histories from finite-locus sampling variation:

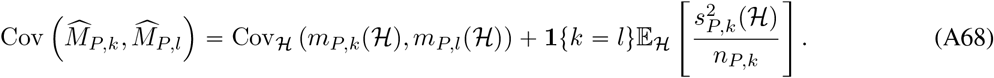

The first term is generally non-diagonal because all five bins share one history ( ℋ) in a Monte Carlo replicate. The second term is diagonal under independent finite-locus sampling across disjoint bins.

Numerically, we discretized the generator in equation (A58) as a nearest-neighbor continuous-time Markov chain on 101 equally spaced frequency states. The boundary rates retain symmetric mutation and conserve total probability. We used implicit time steps of at most 100 generations, 12 generalized Gauss–Laguerre nodes for (*α*^*j*^) ^2^, 200 shared-background histories per *V*_*S*_, and 5,000 bootstrap refits. The finite-locus component used *n*_*P,k*_ *=* 100,000 in every bin as a proxy for the number of variants contributing to the latent S-LDSC target. It does not simulate uncertainty in the S-LDSC regression coefficients themselves, so its uncertainty estimate should be regarded as a lower bound on the actual uncertainty.

**Figure A1:**
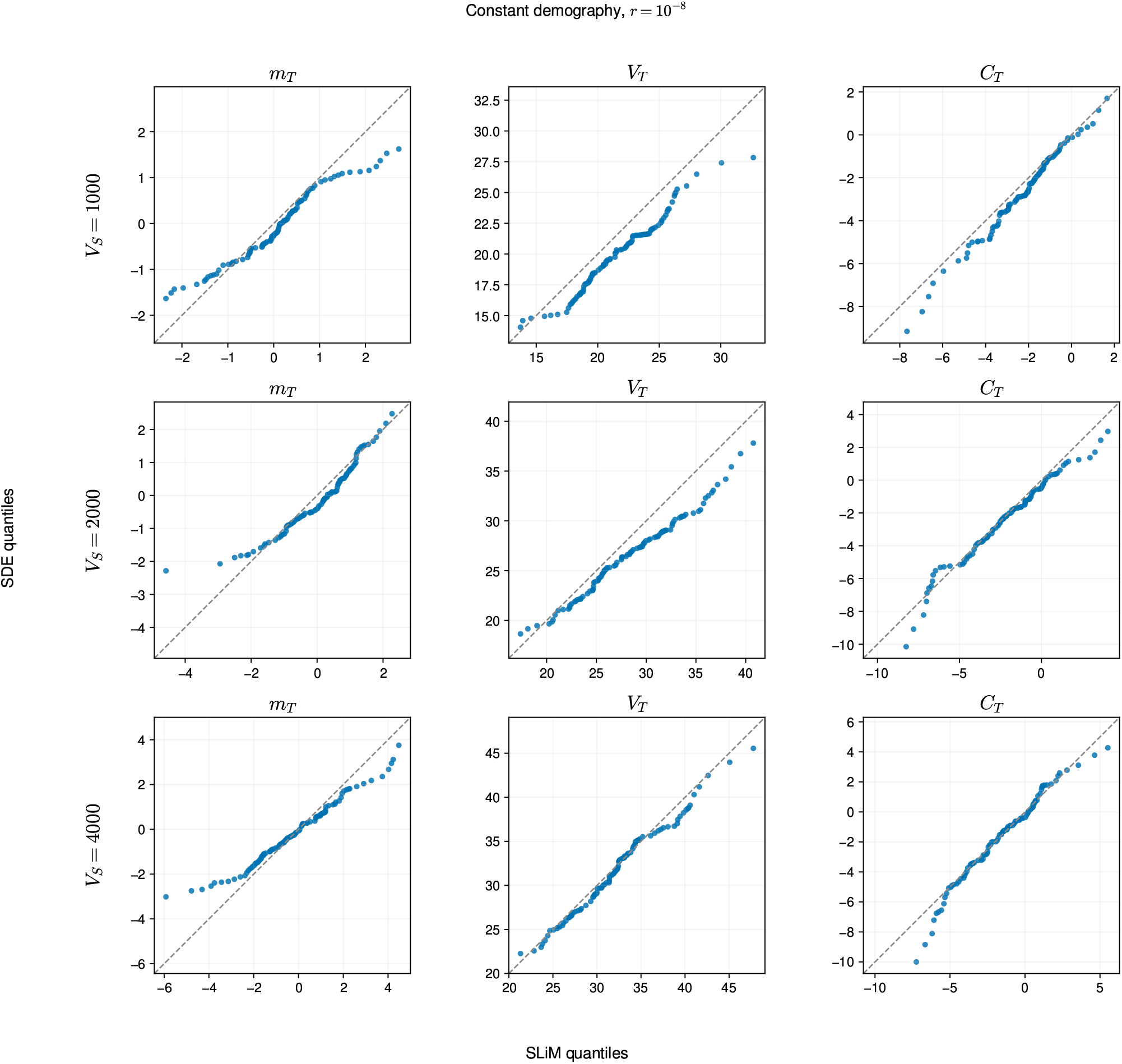
Quantile-Quantile plot of population mean *m*_*t*_, genic variance *V*_*t*_, and LD *C*_*t*_ at generation 8,000 for a diploid population of size *N*_*e*_ *=* 1, 000, comparing SLiM and SDE simulations. We used mutation rate *u =* 10^−10^, recombination rate *r =* 1 × 10^−8^, genome length 1 × 10^8^, and mutational variance 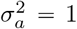. Before computing the harmonic mean, each pairwise recombination fraction was replaced with *c*_*ij*_ + 1 / (2*N*_*e*_ ) − *c*_*ij*_ */* (2*N*_*e*_) to account for finite population size.

**Figure A2:**
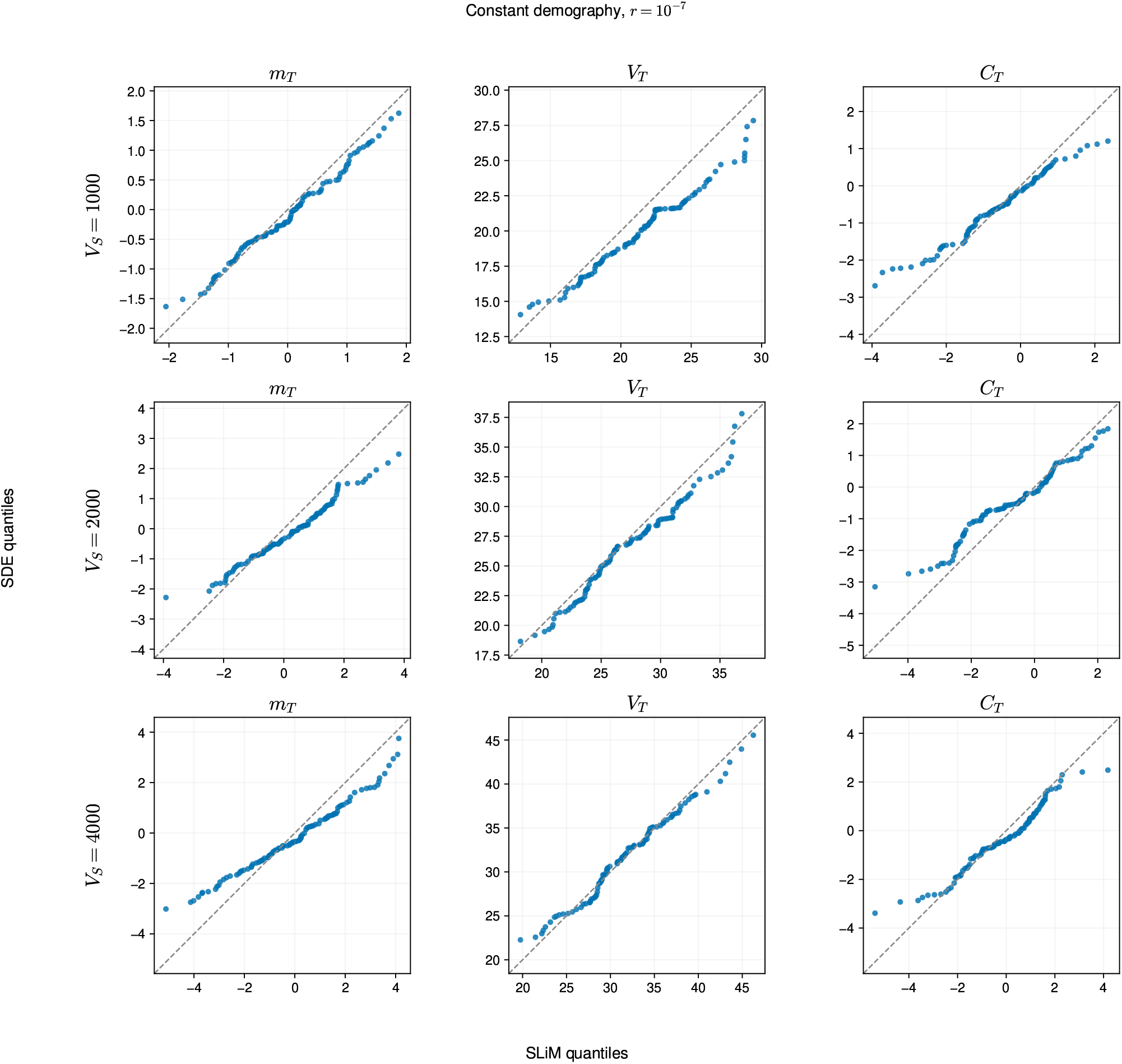
Quantile-Quantile plot of population mean *m*_*t*_, genic variance *V*_*t*_, and LD *C*_*t*_ at generation 8,000 of a diploid population of size *N*_*e*_ 1, 000 comparing SLiM and SDE simulations. We used mutation rate *u* 10^−10^, recombination rate *r =* 1 × 10^−7^, genome length 1 × 10^8^, and mutational variance 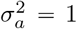. Before computing the harmonic mean, each pairwise recombination fraction was replaced with *c*_*ij*_ 1 /( 2*N*_*e*_) − *c*_*ij*_ */*(2*N*_*e*_) to account for finite population size.

**Figure A3:**
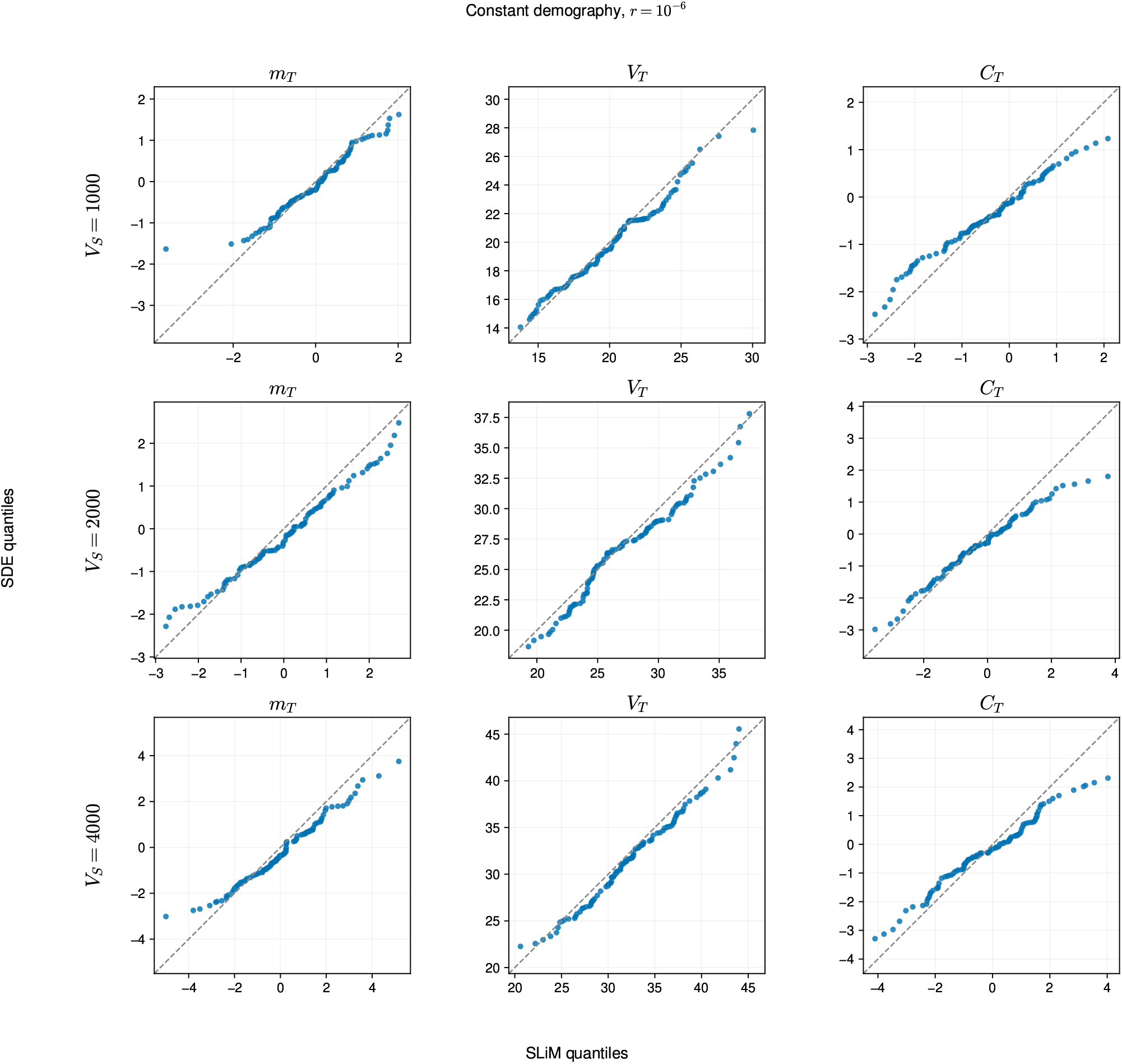
Quantile-Quantile plot of population mean *m*_*t*_, genic variance *V*_*t*_, and LD *C*_*t*_ at generation 8,000 of a diploid population of size *N*_*e*_ *=* 1, 000 comparing SLiM and SDE simulations. We used mutation rate *u* 10^−10^, recombination rate *r =* 1 × 10^−6^, genome length 1 10^8^, and mutational variance *σ*^2^ 1. Before computing the harmonic mean, each pairwise recombination fraction was replaced with *c*_*ij*_ + 1 /(2*N*_*e*_) − *c*_*ij*_ */* (2*N*_*e*_) to account for finite population size.

**Figure A4:**
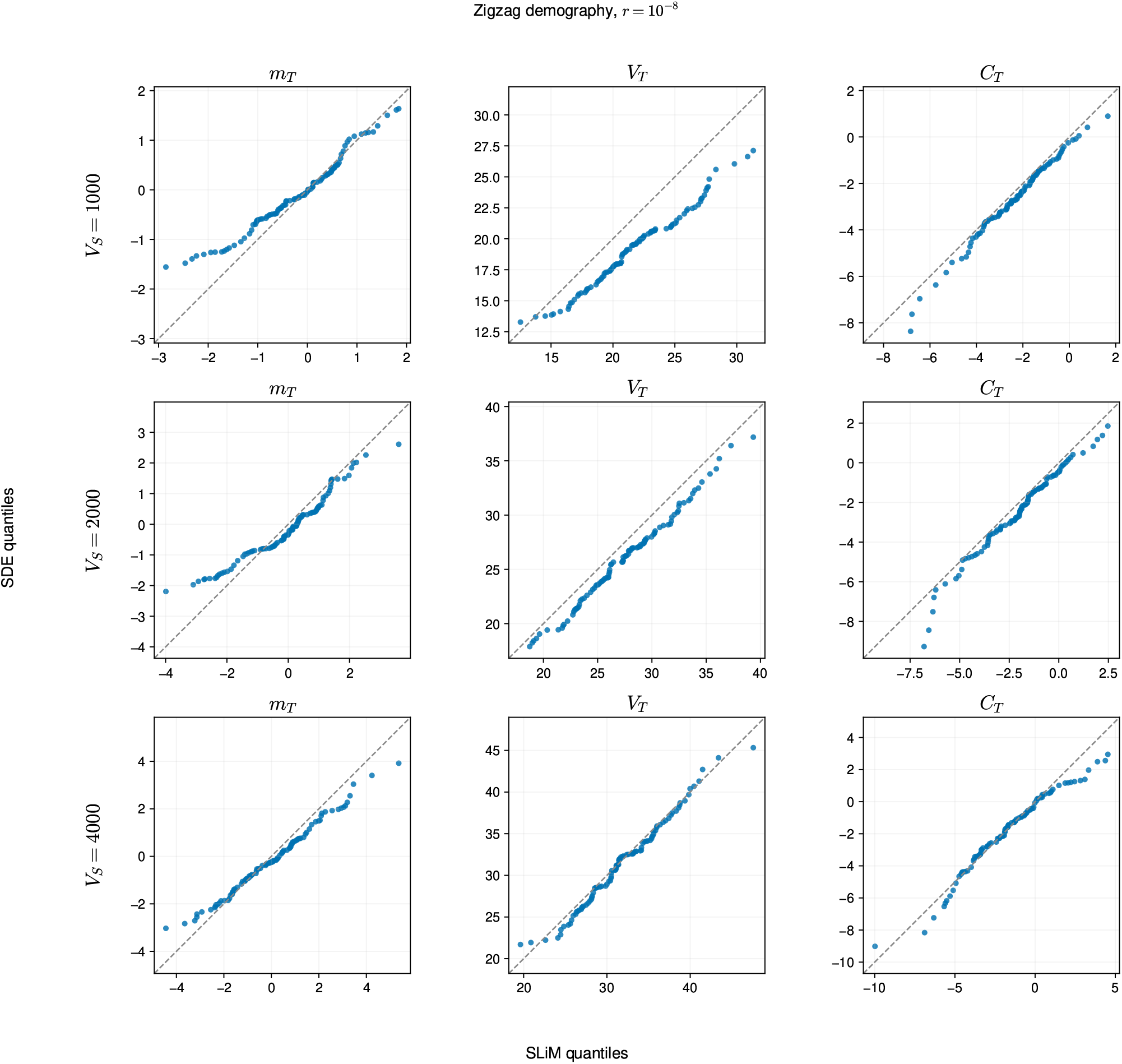
Quantile–quantile plots comparing the terminal distributions of population mean *m*_*t*_, genic variance *V*_*t*_, and LD *C*_*t*_ from SLiM and SDE simulations under changing demography at recombination rate *r* = 10^−8^. The modified `Zigzag_1S14` history (Schiffels and Durbin, 2014) contains three expansions and two declines, alternates exponentially between *N*_*e*_ = 500 and *N*_*e*_ = 2,000, and ends at generation 8,000. Rows correspond to *V*_*S*_ = 1,000, 2,000, and 4,000.

**Figure A5:**
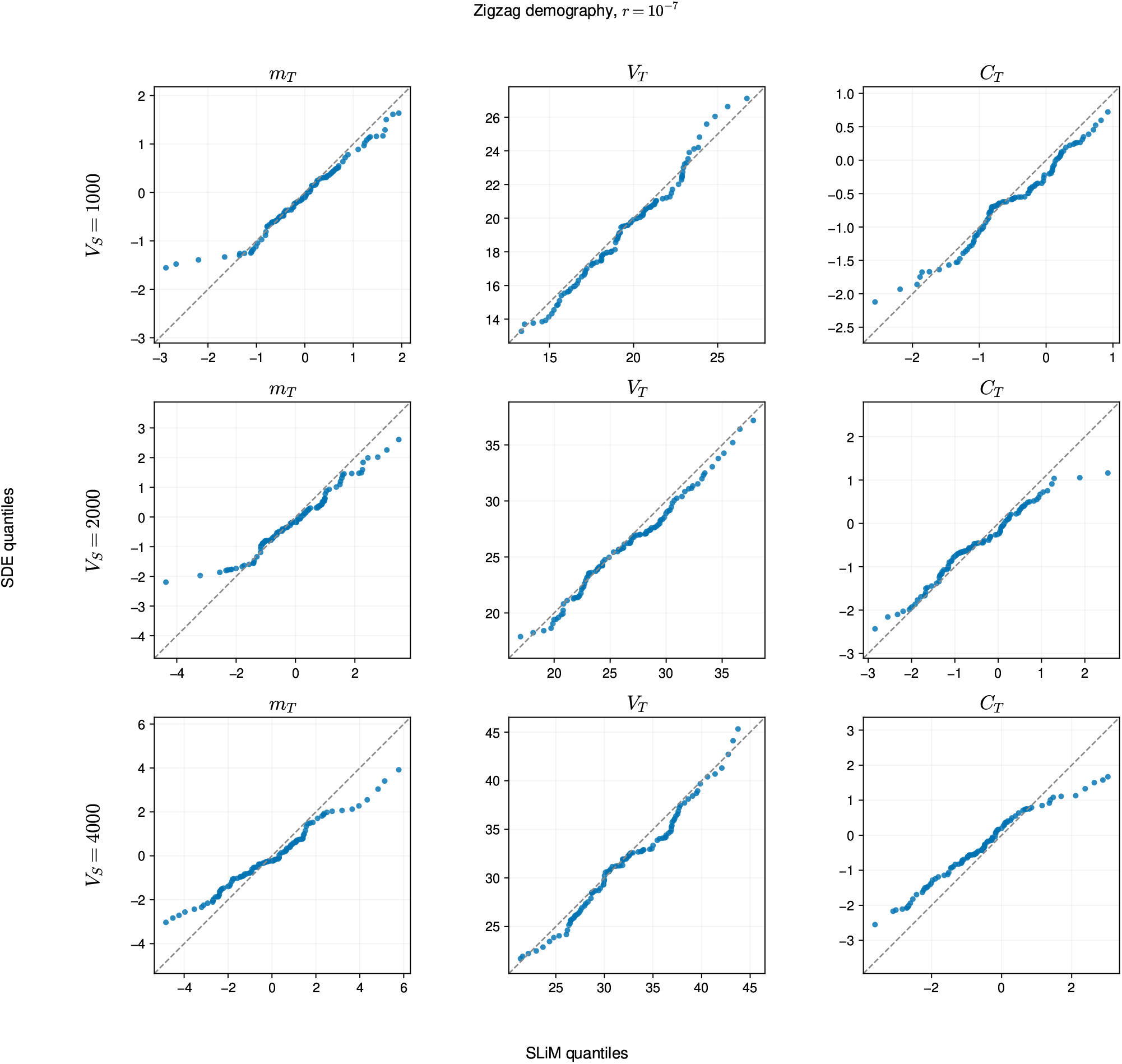
Quantile–quantile plots comparing the terminal distributions of population mean *m*_*t*_, genic variance *V*_*t*_, and LD *C*_*t*_ from SLiM and SDE simulations under the modified `Zigzag_1S14` demography at recombination rate *r* = 10^−7^. Population size changes exponentially between *N*_*e*_ = 500 and *N*_*e*_ = 2,000 across three expansions and two declines; rows correspond to *V*_*S*_ = 1,000, 2,000, and 4,000.

**Figure A6:**
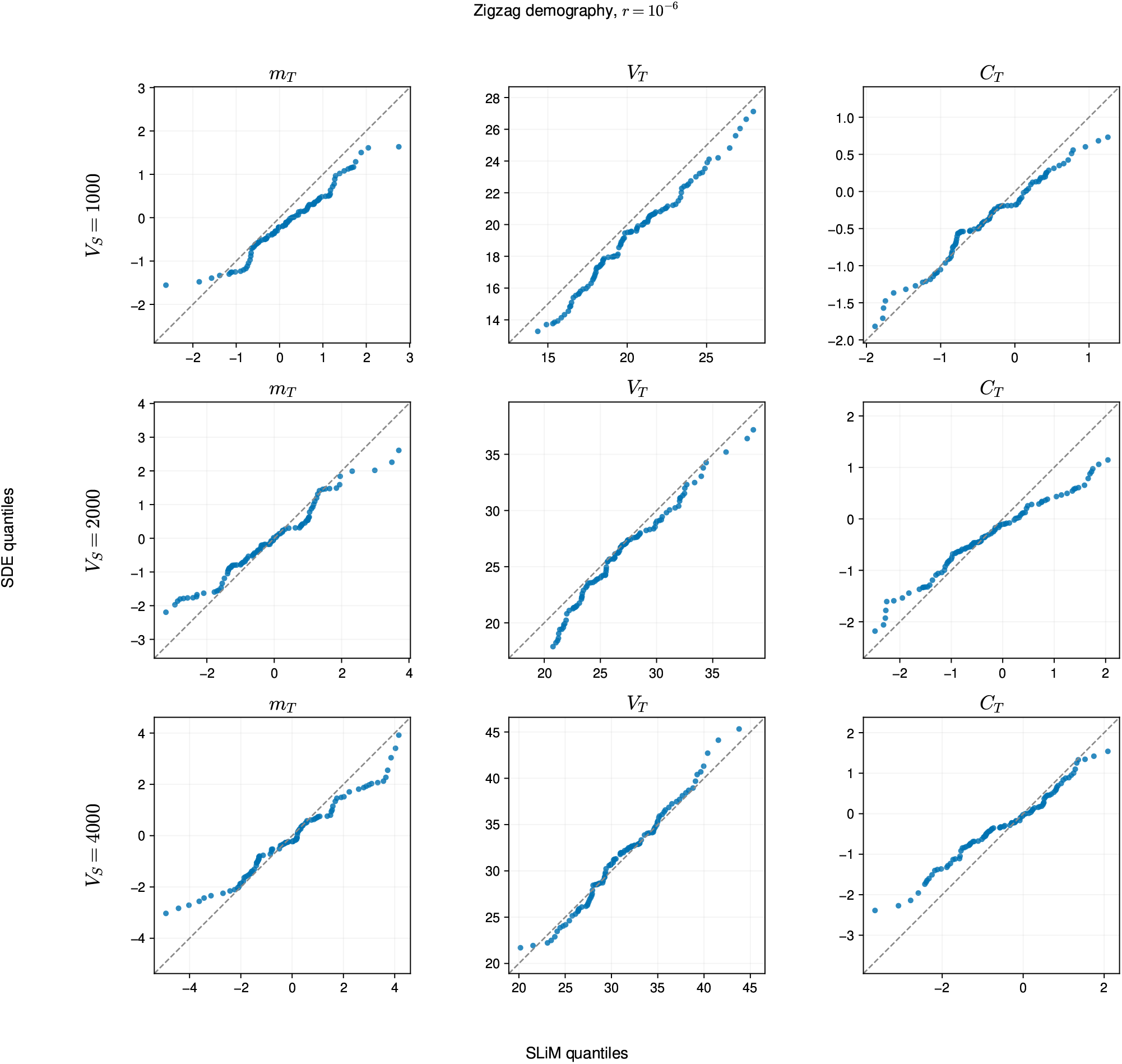
Quantile–quantile plots comparing the terminal distributions of population mean *m*_*t*_, genic variance *V*_*t*_, and LD *C*_*t*_ from SLiM and SDE simulations under the modified `Zigzag_1S14` demography at recombination rate *r* = 10^−6^. Population size changes exponentially between *N*_*e*_ = 500 and *N*_*e*_ = 2,000 across three expansions and two declines; rows correspond to *V*_*S*_ = 1,000, 2,000, and 4,000.

